# Antibodies raised against a structurally defined Aβ oligomer mimic protect human iPSC neurons from Aβ toxicity at sub-stoichiometric concentrations

**DOI:** 10.1101/2025.05.16.654602

**Authors:** Sarah M. Ruttenberg, Rakia Dhaoui, Adam G. Kreutzer, James S. Nowick

## Abstract

Anti-Aβ antibodies are important tools for identifying structural features of aggregates of the Aβ peptide and are used in many aspects of Alzheimer’s disease (AD) research. Our laboratory recently reported the generation of a polyclonal antibody, pAb_2AT-L_, that is moderately selective for oligomeric Aβ over monomeric and fibrillar Aβ and recognizes the diffuse peripheries of Aβ plaques in AD brain tissue but does not recognize the dense fibrillar plaque cores. This antibody was generated against 2AT-L, a structurally defined Aβ oligomer mimic composed of three Aβ-derived β-hairpins arranged in a triangular fashion and covalently stabilized with three disulfide bonds. In the current study, we set out to determine if pAb_2AT-L_ is neuroprotective against toxic aggregates of Aβ and found that pAb_2AT-L_ protects human iPSC-derived neurons from Aβ_42_-mediated toxicity at molar ratios as low as 1:100 antibody to Aβ_42_, with a ratio of 1:25 almost completely rescuing cell viability. Few other antibodies have been reported to exhibit neuroprotective effects at such low ratios of antibody to Aβ. ThT and TEM studies indicate that pAb_2AT-L_ delays but does not completely inhibit Aβ_42_ fibrillization at sub-stoichiometric ratios. The ability of pAb_2AT-L_ to inhibit Aβ_42_ toxicity and aggregation at sub-stoichiometric ratios suggests that pAb_2AT-L_ binds toxic Aβ_42_ oligomers and does not simply sequester monomeric Aβ_42_. These results further suggest that toxic oligomers of Aβ_42_ share significant structural similarities with 2AT-L.

**TOC Graphic:** 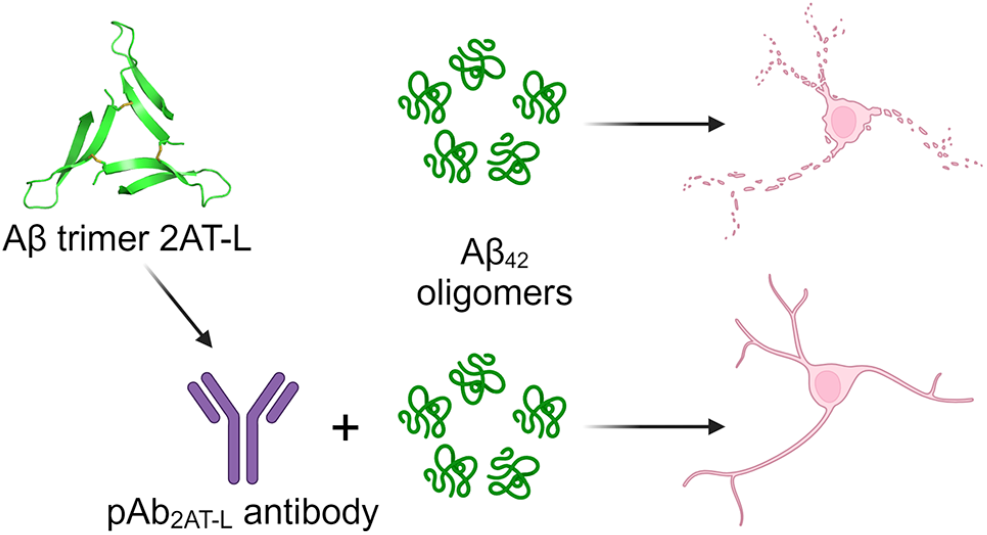

## Introduction

Research over the past two decades has identified oligomers as the most toxic form of Aβ [1-6]. Aβ oligomers contribute to Alzheimer’s disease (AD) pathogenesis by causing neurodegeneration and inflammation in the brain [1-3,6]. Identifying and targeting these species in AD has proven difficult because of the heterogeneity and metastability of Aβ oligomer populations and the low abundance of oligomers in AD pathology compared to monomeric and fibrillar Aβ [1-5,7,8]. Although methods exist to isolate oligomers from AD brain tissue, these methods cannot provide high-resolution structural data and can potentially alter the structures of the oligomers during the isolation or characterization processes [1,2,5,8]. No high-resolution structures of Aβ oligomers isolated from AD brain tissue have been elucidated. Techniques have emerged to stabilize Aβ oligomers formed *in vitro*, some of which have provided high-resolution structures, but the biological relevance of these structures remains to be determined [4,5,7,9,10]. Aβ oligomers can exhibit a variety of morphologies and aggregation states, some of which ultimately form Aβ fibrils like those observed in the brains of AD patients, but tendency to fibrillize does not necessarily correlate with toxicity [1-5,7].

Anti-Aβ antibodies are valuable tools for determining the relevance of various Aβ oligomers in AD pathology and are used in many aspects of AD research. Hundreds of antibodies have been generated against various fragments and species of Aβ. Some target linear epitopes - specific residues of Aβ, while others target conformational epitopes present in oligomeric or fibrillar Aβ which may or may not be residue specific as well. These antibodies have been used for biomarker detection, characterization or isolation of Aβ aggregates, studying the relationship between Aβ structure and toxicity, and a few have been approved for use in AD therapeutics research [1-5,11,12]. Although some antibodies have been developed that target oligomeric Aβ, the epitopes of most of these antibodies are structurally undefined. Antibodies developed against homogenous, structurally defined Aβ oligomers can provide stronger evidence for the relevance of certain Aβ oligomer structures in AD pathogenesis [2-4,6,11].

Our laboratory has synthesized covalently stabilized Aβ oligomer mimics, elucidated their structures at high-resolution, and used them to develop antibodies [4,10,13,14]. These oligomer mimics are composed of peptides designed to mimic β-hairpin conformations in toxic Aβ oligomers. We recently reported the high-resolution structure of one of our Aβ oligomer mimics, 2AT-L (Fig 1). 2AT-L is a toxic trimer composed of three β-hairpins derived from Aβ arranged in a triangular fashion and covalently stabilized with three disulfide bonds [14]. We also reported the generation and study of a polyclonal antibody raised against 2AT-L, pAb_2AT-L_. pAb_2AT-L_ is moderately selective for oligomeric Aβ over monomeric and fibrillar Aβ and stains the diffuse peripheries of Aβ plaques in Down syndrome AD human brain tissue and 5xFAD transgenic mouse brain tissue but does not bind the dense fibrillar plaque cores. The staining of pAb_2AT-L_ in AD tissue suggests that 2AT-L shares structural similarities with assemblies of Aβ present in AD pathology.

**Figure 1.**
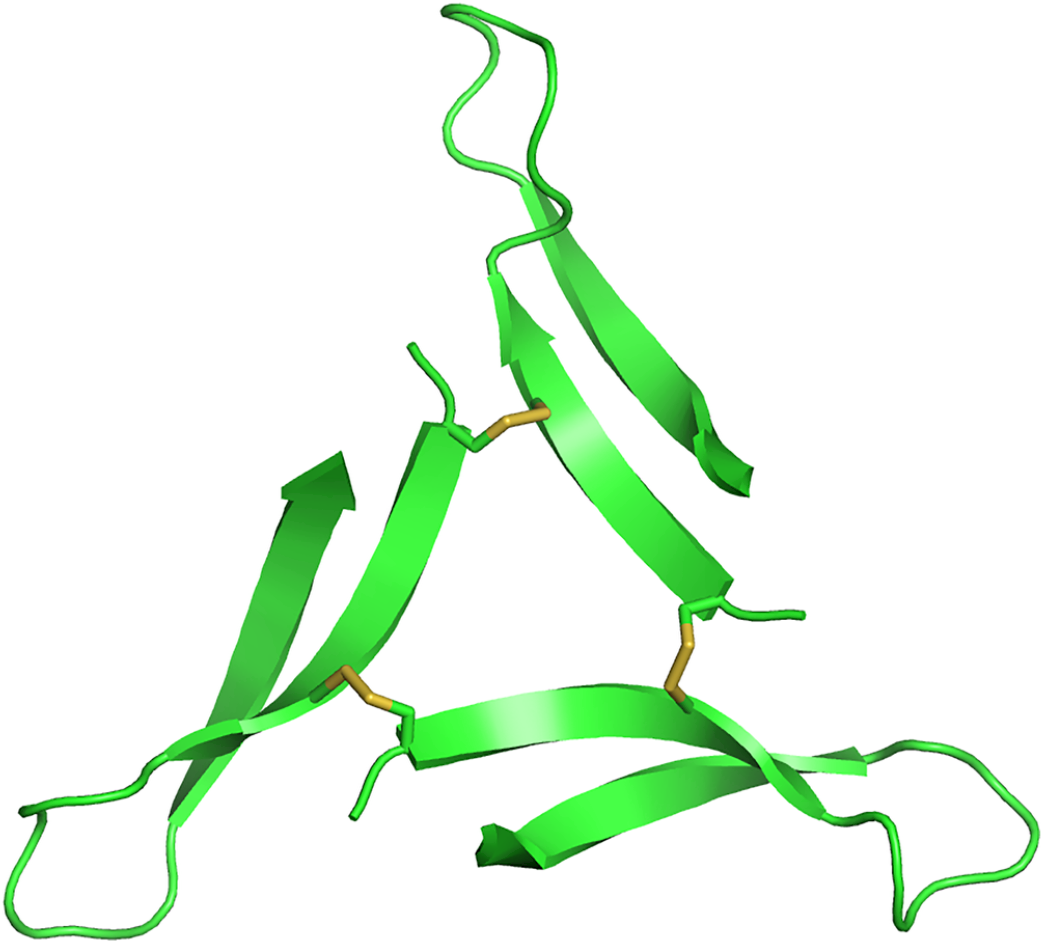
X-ray crystallographic structure of the Aβ oligomer mimic, 2AT-L (PDB 7U4P).

In the current study, we set out to determine if pAb_2AT-L_ is neuroprotective against toxic aggregates of Aβ. We chose to use human iPSC-derived neurons for these experiments because they are a better model for the human neurons affected by AD than immortalized mammalian cell lines or primary rodent neurons and are a valuable research tool because living human neurons cannot be harvested for research [15]. iPSC-derived neurons are neurons created from human stem cells that have been treated to induce neuronal differentiation. We thus examined the effects of Aβ_42_ on iPSC-derived neurons in the presence of various concentrations of pAb_2AT-L_. We further studied the ability of pAb_2AT-L_ to mitigate the production of pro-inflammatory cytokines induced by Aβ_42_ in an immortalized human-derived microglia cell line, HMC3. Finally, we examined the effects of pAb_2AT-L_ on Aβ_42_ aggregation through thioflavin T (ThT) fluorescence assays and transmission electron microscopy (TEM).

## Results

### pAb_2AT-L_ protects human iPSC-derived neurons from Aβ_42_-mediated toxicity

Human iPSC-derived neurons are increasingly being used to study neurodegenerative diseases. Simple methods for generating iPSC-derived neurons have recently emerged, enabling the use of human neurons in research [16]. To investigate whether pAb_2AT-L_ can mitigate the neurotoxicity of Aβ, we studied the effect of recombinant Aβ_42_ on i^3^Neurons in the presence and absence of pAb_2AT-L_. i^3^Neurons (i^3^ = integrated, inducible, and isogenic) are glutamatergic cortical neurons that were differentiated from human iPSCs containing a doxycycline-inducible neurogenin-2 transgene (Ngn2 iPSCs) [17,18]. Treatment with doxycycline causes overexpression of neurogenin-2 which rapidly converts these iPSCs into neurons [17,18]. The Ngn2 iPSCs were designed for use in AD research [18], but to our knowledge, the cytotoxicity of recombinant Aβ toward i^3^Neurons has not been studied.

We generated i^3^Neurons by differentiating Ngn2 iPSCs using the protocols of Ward and coworkers [17] and measured their viability after exposure to Aβ_42_ through two metrics-ATP production (metabolic activity) and LDH release (membrane integrity). In cell-based cytotoxicity assays, the IC50 of Aβ_42_ is typically in the low-micromolar range [19,20,21]. To determine if Aβ_42_ is cytotoxic to i^3^Neurons, we first treated the cells with recombinant Aβ_42_ at concentrations ranging from 25 μM to 98 nM and assayed for viability after 48 hours. Treatment with Aβ_42_ reduced ATP production and increased LDH release in a concentration-dependent manner, with 25 μM Aβ_42_ completely preventing measurable ATP production (Fig 2). Generally, concentrations of Aβ_42_ above 3 μM caused significant reduction in ATP production, increase in LDH release, and morphological changes in the i^3^Neurons. We subsequently determined that, at this concentration of Aβ_42_, a greater degree of LDH release occurred by 72 hours than by 48 hours.

**Figure 2.**
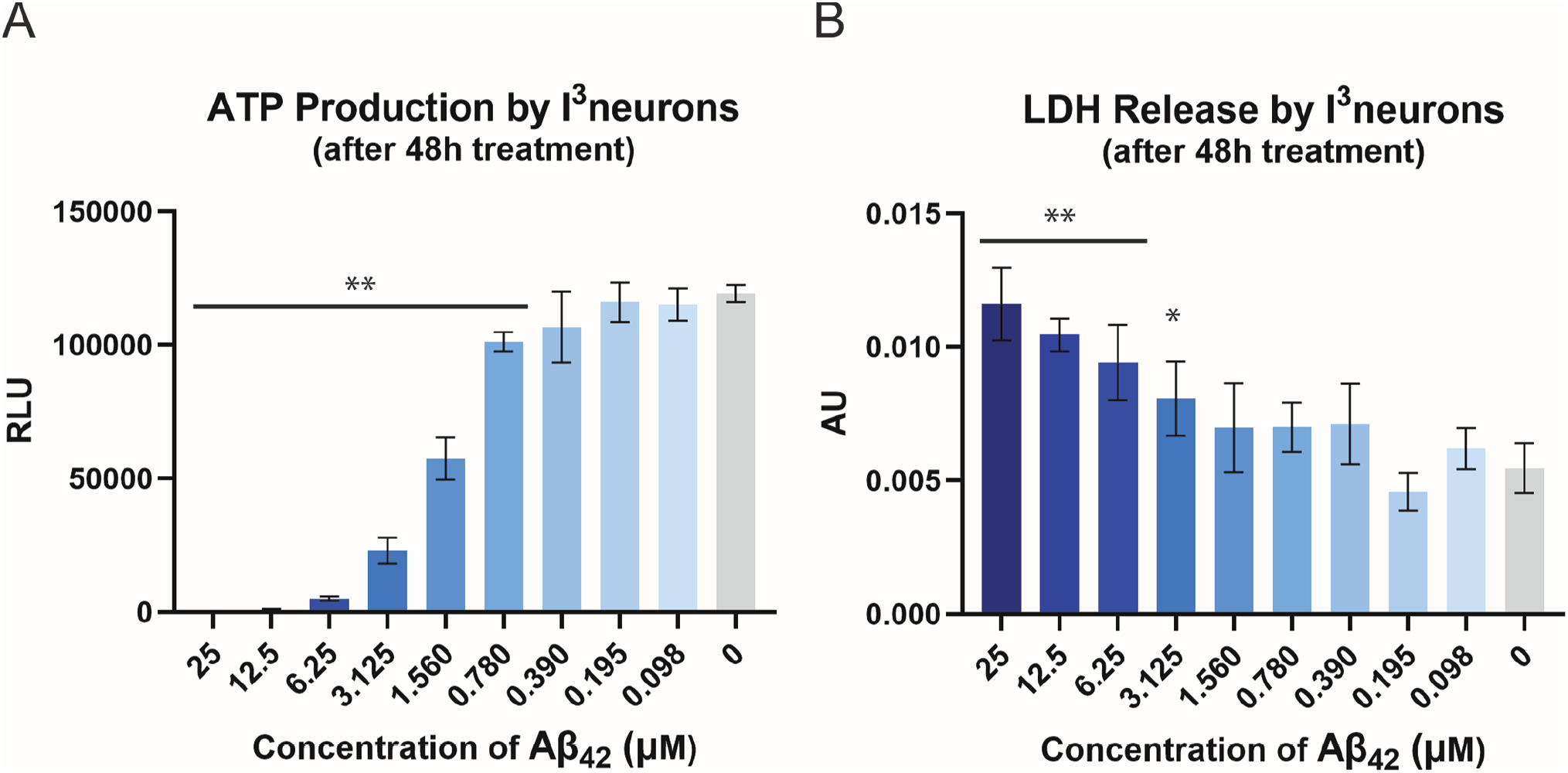
Aβ42 toxicity in i3Neurons. Graphs of (A) ATP production and (B) LDH release by i3Neurons in the presence of varying concentrations of Aβ42 (n=6 technical replicates). ** Significantly different (p < 0.01) from i3neurons treated with PBS (vehicle for AB42).

For subsequent assays, i^3^Neurons were treated with 5 μM Aβ_42_ and pAb_2AT-L_ at concentrations ranging from 200 to 6 nM in triplicate for 72 hours. A positive control of 5 μM Aβ_42_ without pAb_2AT-L_ and a negative control of phosphate buffered saline pH 7.4 (PBS) were also included. After 72 hours of treatment, we measured the relative amounts of ATP and LDH produced by the neurons. pAb_2AT-L_ exhibited significant protective effects on the i^3^Neurons at or above 50 nM -a 1:100 molar ratio of antibody to Aβ_42_ (Fig 3). The protective effects of pAb_2AT-L_ were concentration dependent with 200 nM (a 1:25 molar ratio of antibody to Aβ_42_) almost completely rescuing the cells from Aβ_42_-mediated toxicity (Fig 3). ATP production was restored to levels consistent with the negative control at 200 nM pAb_2AT-L_ (Fig 3A). LDH release was reduced to levels just above the negative control at 200 nM pAb_2AT-L_ (Fig 3B).

**Figure 3.**
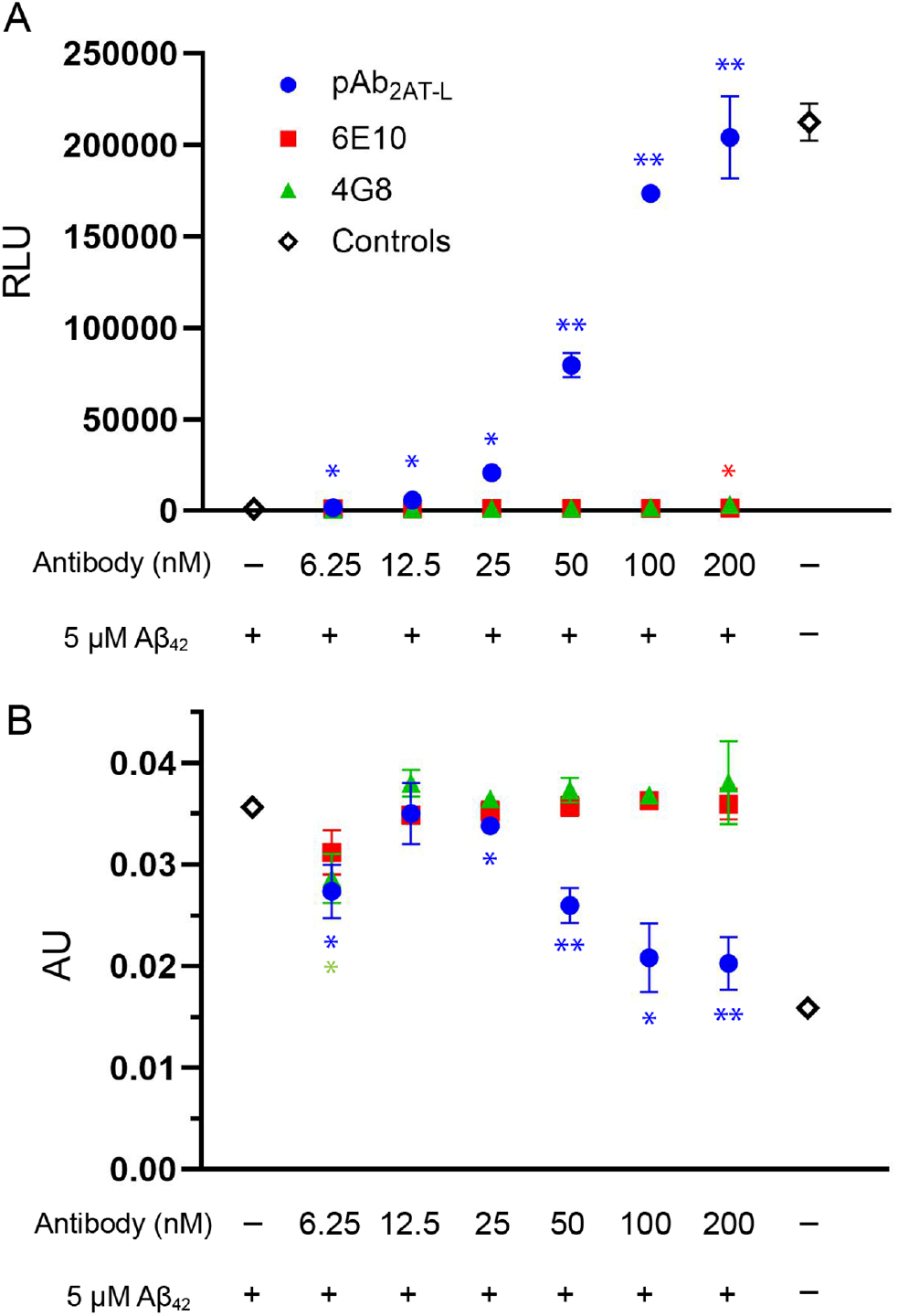
pAb2AT-L protects i3Neurons from Aβ42-mediated toxicity as measured by ATP production and LDH release. (A) ATP production by i3Neurons in the presence of Aβ42 with varying concentrations of pAb2AT-L, measured by the Promega CellTiter-Glo 2.0 assay (n=3 t technical replicates). (B) LDH release by i^3^Neurons in the presence of Aβ42 with varying concentrations of pAb2AT-L, measured by the CyQuant LDH assay (n=3 technical replicates). ** Significantly different (*p* < 0.01) from i^3^neurons treated with just Aβ42. *Significantly different (*p* < 0.05) from i^3^neurons treated with just Aβ_42_.

We compared the protective effects of pAb_2AT-L_ to those of the commercially available anti-Aβ antibodies 6E10 and 4G8; these antibodies are the most widely used in AD research [22]. 6E10 is a monoclonal antibody that targets residues 1-16 of Aβ, and 4G8 is a monoclonal antibody that targets residues 17-24 of Aβ [23]. 6E10 and 4G8 have been shown to bind monomeric, oligomeric, and fibrillar Aβ, with 6E10 having some preference for monomeric and oligomeric over fibrillar. 6E10 has previously been shown to disaggregate Aβ fibrils and increase Aβ neurotoxicity against SH-SY5Y neuroblastoma cells [24]. In the same study, 4G8 had no effect on cell viability or the aggregation state of Aβ [24]. To our knowledge, the protective effects of either of these antibodies against Aβ_42_ have not been studied in iPSC-derived neurons.

In i^3^Neurons treated with 5uM Aβ_42_, 6E10 and 4G8 did not protect against the neurotoxic effects of Aβ_42_ at the concentrations tested. ATP production was not restored, nor were the levels of LDH reduced at any concentrations tested of these antibodies (200-6.25 nM). This observation is consistent with previous studies of 6E10 and 4G8 in SH-SY5Y cells [24,25]. To confirm that the protective effects exhibited by pAb_2AT-L_ were not a general effect of rabbit IgG protein, we performed the same experiment with a generic rabbit IgG antibody. The generic rabbit IgG antibody did not have any protective effects against Aβ_42_ in i^3^Neurons (S1 Fig.).

To study the protective effect of pAb_2AT-L_ further, we also tested the ability of pAb_2AT-L_ to inhibit Aβ_42_-induced pro-inflammatory cytokine production by HMC3 microglia. Treatment with Aβ_42_ causes HMC3 microglia to produce the pro-inflammatory cytokine IL-6 (S2 Fig.) [26-28]. IL-6 is also upregulated in AD animal models and human AD patients [29]. The two highest concentrations of pAb_2AT-L_ tested, 200 nM and 100 nM, significantly reduced Aβ_42_-induced IL-6 production in HMC3 microglia (*p* < 0.01) (S2 Fig.). At 200 nM pAb_2AT-L_, IL-6 production was reduced by about 50% compared to the controls. This result corroborates that pAb_2AT-L_ sequesters toxic Aβ_42_ species and protects cells at sub-stoichiometric concentrations.

### ThT assay of Aβ42 with pAb_2AT-L_

To better understand the interactions between pAb_2AT-L_ and Aβ_42_, we investigated the effects of pAb_2AT-L_ on Aβ_42_ aggregation. First, we monitored the fibrilization of a 3 μM solution of recombinant Aβ_42_ using thioflavin T in the presence and absence of pAb_2AT-L_. In the absence of pAb_2AT-L_, the ThT signal began to increase after one hour, plateauing after 2 hours (Fig 4). In the presence of pAb_2AT-L_, Aβ_42_ fibrillization occurred more slowly. At the lowest concentration of pAb_2AT-L_, 210 nM, Aβ_42_ fibrillization was delayed by about an hour. Increasing concentrations of pAb_2AT-L_ led to increasing delays in Aβ_42_ fibrillization. At 800 nM pAb_2AT-L_, ThT-positive aggregates of Aβ_42_ did not appear to form until after five hours, indicating that pAb_2AT-L_ delayed fibrillization by about four hours.

**Figure 4.**
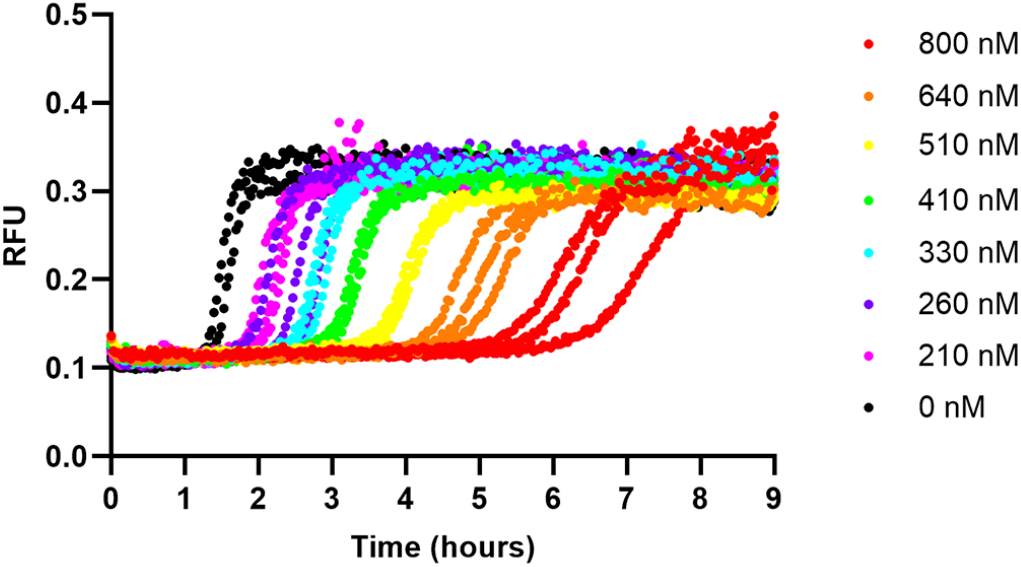
pAb2AT-L delays fibril formation of Aβ42. ThT fluorescence assay of 3 μM Aβ42 in the presence of varying concentrations of pAb2AT-L (210-800 nM).

### TEM of Aβ42 with pAb_2AT-L_

We performed transmission electron microscopy (TEM) to visualize the effect of pAb_2AT-L_ on Aβ_42_ aggregation. To do this, we prepared a 3 μM solution of lyophilized recombinant Aβ_42_ dissolved in PBS and incubated in the presence or absence of 800 nM pAb_2AT-L_ for four hours. We then applied the solutions to TEM grids, stained with a 1 % uranyl acetate solution, and imaged the grids. In the absence of pAb_2AT-L_, Aβ_42_ formed bundles of fibrillar structures (Fig 5). The protofibrils making up the bundles appear to be approximately 200-1000 nm in length and were often observed in the presence of spherical or amorphous aggregates. In the presence of pAb_2AT-L_ under the same conditions, Aβ_42_ did not appear to form fibrillar structures (Fig 5). Instead, only amorphous aggregates were observed. We also applied a solution of pAb_2AT-L_ to TEM grids and observed solids. Collectively, these TEM studies corroborate the findings from the ThT studies that sub-stoichiometric pAb_2AT-L_ inhibits Aβ_42_ fibrillization.

**Figure 5.**
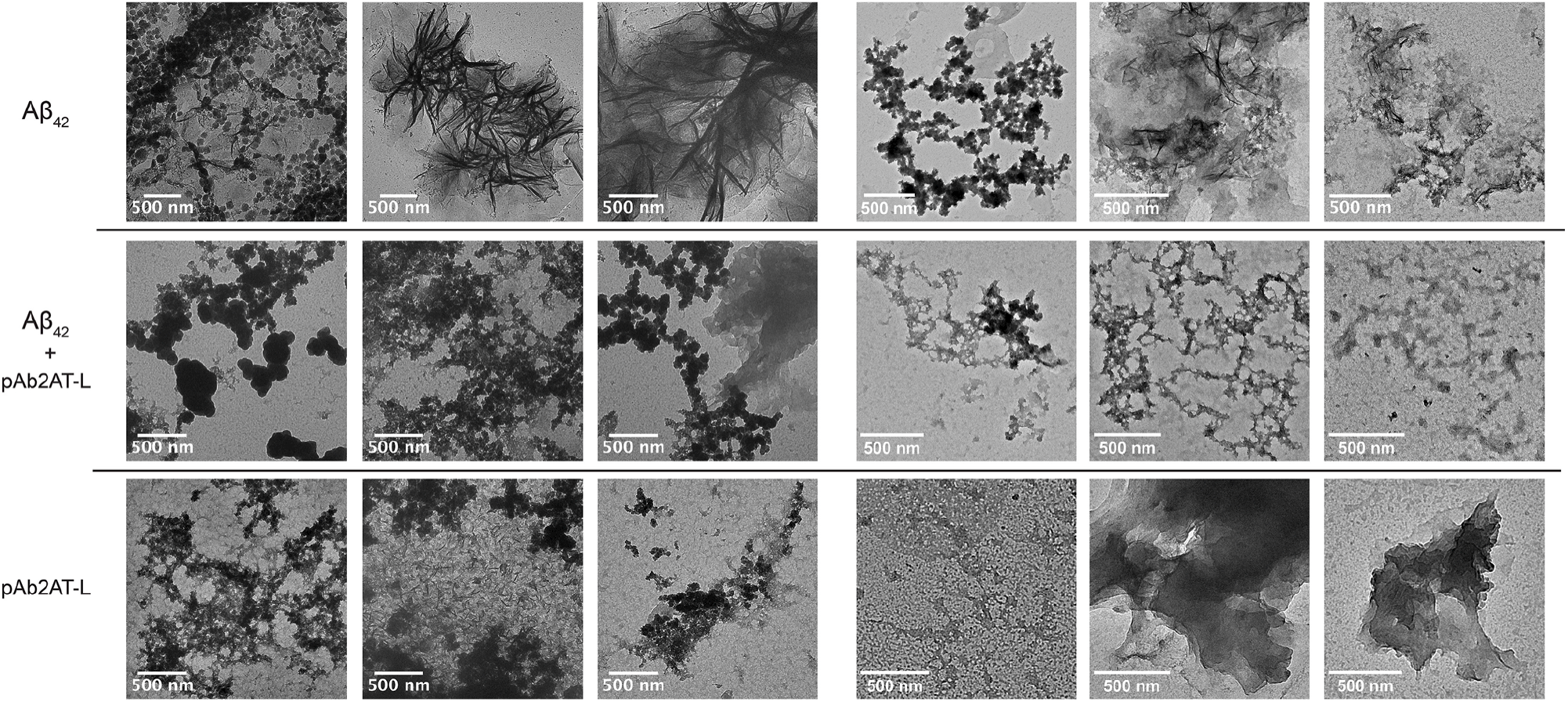
pAb2AT-L inhibits Aβ42 fibril formation as visualized by TEM imaging. Representative TEM images of 3 uM Aβ42 (Top), 3 uM Aβ42 and 800 nM pAb2AT-L (middle), 800 nM pAb2AT-L (bottom) from two separate TEM sessions (left and right). After four hours of incu in PBS, samples were applied to carbon mesh copper grids and stained with uranyl acetate (UA) before imaging (the session on the left was stained with 2 % UA and the session on the right was stained with 1 % UA).

## Discussion

A11 and OC are the most commonly used polyclonal anti-Aβ antibodies in AD research. While OC binds fibrils and fibrillar oligomers of Aβ, A11 binds high-molecular weight (∼40 kDa or larger) non-fibrillar oligomers of Aβ, as well as a variety of other amyloid oligomers [30,31]. Glabe and coworkers originally generated A11 against Aβ_40_ oligomers conjugated to gold nanoparticles [30]. At equimolar concentrations or more, A11 protects SH-SY5Y cells from the toxicity of A11-positive Aβ_40_ and Aβ_42_ oligomers, but at lower concentrations A11 exhibits minimal inhibition of Aβ oligomer-mediated toxicity [30,32,33]. It is noteworthy that pAb_2AT-L_ at nanomolar concentrations substantially alters the toxicity and aggregation of Aβ at micromolar concentrations. There are few reports of antibodies affording significant neuroprotective effects at such low ratios of antibody to Aβ [34-42]. Lecanemab, a monoclonal anti-Aβ antibody that preferentially binds protofibrils of Aβ, has been shown to completely rescue PC12 cells from the toxic effects of Aβ_42_ protofibrils at sub-stoichiometric ratios. For 1 uM Aβ_42_ protofibrils, the ED50 of Lecanemab was 13 nM [40]. Additionally, some Aβ oligomer-specific antibodies and antibody fragments exhibit protective effects against toxic Aβ oligomers at sub-stoichiometric ratios [34-39,41]. The most protective of these exhibited protective effects on SH-SY5Y cells at ratios of antibody to Aβ_42_ oligomers as low as 1:100 in some cases [36]. Most other anti-Aβ antibodies have not been reported to exhibit substantial protective effects on cells at less than equimolar concentrations of antibody to Aβ_42_[33,43-50].

The sub-stoichiometric activity of pAb_2AT-L_ against the toxicity and aggregation of Aβ_42_ suggests that pAb_2AT-L_ can bind epitopes displayed on toxic Aβ_42_ oligomers and is not simply sequestering monomeric Aβ_42_ and preventing its aggregation. If pAb_2AT-L_ were only binding monomeric Aβ_42_, only a small fraction of Aβ_42_ would be sequestered at the concentrations of pAb_2AT-L_ that were tested, and this would not result in the significant reduction of Aβ_42_-mediated toxicity. Instead, it appears that pAb_2AT-L_ must be binding non-fibrillar aggregates (oligomers) of Aβ_42_. Although the data presented here do not definitively confirm that pAb_2AT-L_ does not bind fibrillar Aβ_42_, the ThT and T data show that pAb_2AT-L_ delays Aβ_42_ fibrilli ation which suggests that pAb_2AT-L_ binds Aβ_42_ species before fibril formation. One of the limitations of the cell-based experiments is that during the 72 hours in which Aβ_42_ is incubated with the i^3^Neurons, a variety of aggregates form and thus we cannot be certain of the exact toxic species. Nevertheless, we previously showed that pAb_2AT-L_ has a moderate preference for binding oligomeric Aβ_42_ over monomeric and fibrillar Aβ_42_[14], and the data presented here further support that conclusion.

The unique structure of the 2AT-L antigen used to generate pAb_2AT-L_ is likely responsible for the sub-stoichiometric activity of this antibody. 2AT-L was designed to mimic an Aβ oligomer, specifically a trimer. It lacks the N-terminus of Aβ, presents residues 17-36 in a β-hairpin conformation consisting of residues 17-23 and 30-36 in β-strand conformations and residues 24-29 in a loop conformation. These properties of 2AT-L are likely reflected in some of the epitopes bound by the antibodies present in the polyclonal mixture comprising pAb_2AT-L_. Because pAb_2AT-L_ is a mixture, we cannot be sure which features of the antigen are reflected in the epitopes of the individual antibodies that bind the toxic oligomers formed by Aβ_42_. Among the features that may be reflected is the β-hairpin conformation of the 17-36 segment of Aβ. β-Hairpins are the building blocks of several reported and proposed structures of toxic Aβ oligomers and an affinity for binding Aβ β-hairpin conformations might explain the protective effect afforded by pAb_2AT-L_ [4,7,14,51-57].

## Materials and methods

### Antibody generation

pAb_2AT-L_ was generated in rabbits immunized with 2AT-L conjugated to KLH by Pacific Immunology. The rabbit serum was collected, and affinity purified on an NHS-activated agarose resin column functionalized with 2AT-L.

### Aβ_42_ preparation

Recombinantly expressed Aβ_42_ as the ammonium salt was purchased from rPeptide (catalog# A-1167-2) and received as lyophilized solid. The solid was dissolved in 2 mM NaOH to create a 1 mg/mL Aβ_42_ solution, sonicated for 5 minutes, and aliquoted into 0.02 μmol aliquots. The aliquots were then frozen, lyophilized, and stored at -80° C until use. Solutions of Aβ_42_ were freshly prepared from the lyophilized aliquots for assays.

### i^3^Neuron culture

i^3^Neurons were generated from Ngn2 iPSCs according to the protocols published by Ward and coworkers [17]. Ngn2 iPSCs were obtained from the Blurton-Jones laboratory at the Sue and Bill Gross Stem Cell Research Center. Briefly, Ngn2 iPSCs were thawed and plated in mTeSR media treated with 10 μM rock inhibitor. Cells were plated at a density of 500,000 cells per ml on a Matrigel coated 6-well plate (2 ml per well). Cells were incubated at 37 °C with 5% CO2 and cell media was aspirated and replaced daily with mTesR (without rock inhibitor) until the cell density reached 80% confluency. Cells were then passaged by aspirating the media, treating with accutase, centrifuging, and replating in fresh mTesR with rock inhibitor.

For differentiation, cells were passaged and resuspended in neuronal predifferentiation media (Knockout DMEM:F12, 1x N2, 1x non-essential amino acids, 10 ng/ml BDNF, 10 ng/ml NT3, 1 ug/ml mouse laminin, 2 ug/ml doxyclyine) with rock inhibitor (10 uM), plated in a new Matrigel coated 6-well plate at the same density in 2ml of media per well, and incubated at 37 °C. The following two days media was aspirated and replaced. Cells were then either frozen in Knockout DMEM with 10% DMSO for later maturation or passaged and resuspended in neuronal maturation media (1:1 neurobasal A media and DMEM f:12, 1x B27 0.5x N2, 1x non-essential amino acids, 0.5x glutamax 10 ng/ml BDNF, 10 ng/ml NT3, 1 ug/ml mouse laminin, 2 ug/ml doxyclyine). Cells were plated for maturation on 96-well poly-D-lysine-coated plates at a density of 200,000 cells per ml. After one week of maturation, media was gently removed and replaced with maturation media containing the compounds of interest, but lacking doxycycline. The neurons were incubated for 72 hours before assaying according to manufacturer protocols.

#### HMC3 culture

HMC3 microglia cells were purchased from ATCC (Cat# CRL-3304). Upon arrival, cells were stored at -80 °C for 2 days and then thawed in a bead bath at 37 °C. The cells were diluted in pre-warmed EMEM with 10 % fetal bovine serum and centrifuged. The supernatant was aspirated, and the cell pellet was redissolved in 1 mL of EMEM and transferred to a 75 cm^2^ cell culture flask containing 19 mL of media. The cells were incubated at 37 °C with 5% CO_2_ until reaching 80 % confluence (three to five days) before passaging with trypsin to a new cell culture flask. On the subsequent passage, the cells were plated on 96-well cell culture plates at a density of one hundred thousand cells per mL. incubated overnight before treatment. The cells were incubated for 72 hours before assaying according to manufacturer protocols.

### ThT assay

The ThT assay was performed on 3 μM Aβ_42_ in PBS at pH 7.4 (10 mM Na_2_HPO_4_, 1.8 mM KH_2_PO_4_, 137 mM NaCl, 2.7 mM KCl) containing 10 μM ThT in the presence of a dilution series of pAb_2AT-L_ (800-210 nM, 0nM). The assay was performed in triplicate in Corning® 96-well Half Area Black/Clear Flat Bottom Polystyrene NBS Microplates (product# 3881) at 25 °C under quiescent conditions. Briefly, a serial dilution series of 10x concentrations of pAb_2AT-L_ was prepared in deionized water in a 96-well plate and aliquoted in quadruplicate across four rows of the 96-well plate. A 3.33 μM solution of Aβ_42_ was then prepared in PBS at pH 7.4 (10 mM Na_2_HPO_4_, 1.8 mM KH_2_PO_4_, 137 mM NaCl, 2.7 mM KCl) containing 10 μM thioflavin T (ThT) and added to all but one row of the pAb_2AT-L_ solutions. The plate was sealed with clear adhesive and fluorescence was immediately measured on a ThermoFisher Scientific Varioskan Lux plate reader at excitation/emission wavelengths of 440/485 nm and excitation bandwidth of 12 nm for one second. Measurements were acquired every two minutes for five hours. The data were plotted in GraphPad Prism

### TEM sample preparation and imaging

3 μM solutions of Aβ_42_ were prepared in PBS at pH 7.4 with and without 800 nM pAb_2AT-L_. Samples were incubated for three hours at room temperature without shaking to elicit fibril formation. After incubation, 5uL of sample was deposited on 200-mesh formvar/carbon-coated copper grid carbon-copper grids purchased from Electron Microscopy Sciences (catalog #FCF200-Cu-50) and left to dry for 15 minutes. The grids were then gently wicked, allowed to dry for another 5 minutes, and treated with 5uL of two percent uranyl acetate for 2 minutes for negative staining. The grids were then gently wicked, allowed to dry for 5 minutes, and then washed with 5 μL of nanopure water. After a minute, the water was wicked, and the grids were left to dry for 10 minutes before imaging. Samples were transferred to a JEOL 2100F TEM and imaged using a Schottky type field emission gun operating at 200kV. Images were recorded using a Gatan OneView CMOS camera at 4k x 4k resolution.

## Acknowledgements

The authors thank the Sue and Bill Gross Stem Cell Research Center, specifically Christina Tu for training us to work with iPSCs, and Professor Mathew Blurton-Jones and his laboratory for supplying us with Ngn2 iPSCs and providing guidance for their use. The authors also thank the UC Irvine Materials Research Institute, specifically Li Xing for TEM instrument training and usage.

## Funding statement

This research was funded by the National Institutes of Health (NIH), National Institute on Aging (NIA). J.S.N. was awarded AG072587. The NIA was not involved in this research or manuscript beyond funding. The funders had no role in study design, data collection and analysis, decision to publish, or preparation of the manuscript.

## Competing interest statement

J.S.N and A.G.K are inventors of a patent describing synthetic amyloid oligomers and their applications (U.S. Patent 10,662,226), which is assigned to the UC Regents.

## Supporting Information

This article contains supporting information. Procedures detailing the generation and purification of pAb_2AT-L_, cell culture and differentiation, cell assays, ThT assays, and TEM studies can be found in the supporting information.

## References

1. Cline EN, Bicca MA, Viola KL, Klein WL. The Amyloid-β Oligomer Hypothesis: Beginning of the Third Decade. Journal of Alzheimer’s Disease 2018, 64 (1), S567–S610. doi: 10.3233/jad-179941.

2. Shea D, Daggett V. Amyloid-β Oligomers: Multiple Moving Targets. Biophysica 2022, 2 (2), 91–110. doi: 10.3390/biophysica2020010.

3. Song C, Zhang T, Zhang Y. Conformational Essentials Responsible for Neurotoxicity of Aβ42 Aggregates Revealed by Antibodies against Oligomeric Aβ42. Molecules 2022, 27 (19), 6751. doi: 10.3390/molecules27196751.

4. Ruttenberg SM, Nowick JS. A Turn for the Worse: Aβ β-Hairpins in Alzheimer s Disease. Bioorganic & medicinal chemistry 2024, 105, 117715–117715. doi: 10.1016/j.bmc.2024.117715.

5. Wang Y, Chen J, Gao F, Hu M, Wang, X Recent Developments in the Chemical Biology of Amyloid-β Oligomer Targeting. Organic and Biomolecular Chemistry 2023, 21, 4540–4552. doi: 10.1039/D3OB00509G.

6. Zhang Y, Chen H, Li R, Sterling K, Song W. Amyloid β-Based Therapy for Alzheimer s Disease: Challenges, Successes and Future. Signal Transduction and Targeted Therapy 2023, 8 (1), 1–26. doi: 10.1038/s41392-023-01484-7.

7. Aleksis R, Oleskovs F, Jaudzems K, Pahnke J, Biverstål H. Structural Studies of Amyloid-β Peptides: Unlocking the Mechanism of Aggregation and the Associated Toxicity. Biochimie 2017, 140, 176–192. doi: 10.1016/j.biochi.2017.07.011.

8. Hong W, Wang Z, Liu W, O’Malley TT, Jin M, Willem M, et al. M. Diffusible, Highly Bioactive Oligomers Represent a Critical Minority of Soluble Aβ in Alzheimer’s Disease Brain. Acta Neuropathologica 2018, 36 (1), 19–40. doi: 10.1007/s00401-018-1846-7.

9. Jeon J, Yau WM, Tycko R. Early Events in Amyloid-β Self-Assembly Probed by Time-Resolved Solid-State NMR and Light Scattering. Nature communications 2023, 4 (1). doi: 10.1038/s41467-023-38494-6.

10. Samdin TD, Kreutzer AG, Nowick JS. Exploring Amyloid Oligomers with Peptide Model Systems. Current Opinion in Chemical Biology 2021, 64, 106–115. doi: 10.1016/j.cbpa.2021.05.004.

11. Bitencourt ALB, Campos RM, Cline EN. Klein WL, Sebollela A. Antibody Fragments as Tools for Elucidating Structure-Toxicity Relationships and for Diagnostic/Therapeutic Targeting of Neurotoxic Amyloid Oligomers. International Journal of Molecular Sciences 2020, 2 (23), 8920. doi: 10.3390/ijms21238920.

12. Although some Aβ antibodies have been approved to treat Alzheimer’s disease, their use as a therapeutic remains controversial due to their low efficacy and high incidence of life-threatening side effects.

13. Kreutzer AG, Malonis RJ, Parrocha CM, Tong K, Guaglianone G, Nguyen JT, et al. Generation and Study of Antibodies against Two Triangular Trimers Derived from Aβ. Peptide Science 2023. doi: 10.1002/pep2.24333.

14. Kreutzer AG, Parrocha CM, Haerianardakani S, Guaglianone G, Nguyen J, Diab MN, et al. Antibodies Raised against an Aβ Oligomer Mimic Recognize Pathological Features in Alzheimer’s Disease and Associated Amyloid-Disease Brain Tissue. ACS Central Science 2023. doi: 10.1021/acscentsci.3c00592.

15. Dolmetsch R, Geschwind DH. The Human Brain in a Dish: The Promise of IPSC-Derived Neurons. Cell 2011, 145 (6), 831–834. doi: 10.1016/j.cell.2011.05.034.

16. Cetin S, Knez D, Gobec S, Kos J, Pišlar A. Cell Models for Alzheimer’s and Parkinson’s Disease: At the Interface of Biology and Drug Discovery. Biomedicine & Pharmacotherapy 2022, 149, 112924. doi: 10.1016/j.biopha.2022.112924.

17. Fernandopulle MS, Prestil R, Grunseich C, Wang C, Gan L, Ward ME. Transcription Factor-Mediated Differentiation of Human IPSCs into Neurons. Current Protocols in Cell Biology 2018, 79 (1), e51. doi:10.1002/cpcb.51.

18. Wang C, Ward ME, Chen R, Liu K, Tracy TE, Chen X, et al. Scalable Production of IPSC-Derived Human Neurons to Identify Tau-Lowering Compounds by High-Content Screening. Stem Cell Reports 2017, 9 (4), 1221–1233. doi: 10.1016/j.stemcr.2017.08.019.

19. Raskatov JA. What Is the “Relevant” Amyloid β42 Concentration?. 2019, 20 (13), 1725–1726. doi: 10.1002/cbic.201900097.

20. Dutta S, Foley AR, Warner CJA., Zhang X, Rolandi M, Abrams B, Raskatov JA. Suppression of Oligomer Formation and Formation of Non-Toxic Fibrils upon Addition of Mirror-Image Aβ42 to the Natural L -Enantiomer. Angewandte Chemie 2017, 56 (38), 11506–11510. doi: 10.1002/anie.201706279.

21. Finder VH, Vodopivec I, Nitsch RM, Glockshuber R. The Recombinant Amyloid-β Peptide Aβ1-42 Aggregates Faster and Is More Neurotoxic than Synthetic Aβ1-42. Journal of Molecular Biology 2010, 396 (1), 9–18. doi: 10.1016/j.jmb.2009.12.016.

22. Hunter S, Brayne C. Do Anti-Amyloid Beta Protein Antibody Cross Reactivities Confound Alzheimer Disease Research? Journal of Negative Results in BioMedicine 2017, 16 (1). doi: 10.1186/s12952-017-0066-3.

23. Hatami A, Albay R, Monjazeb S, Milton S, Glabe C. Monoclonal Antibodies against Aβ42 Fibrils Distinguish Multiple Aggregation State Polymorphismsin Vitroand in Alzheimer Disease Brain. Journal of Biological Chemistry 2014, 289 (46), 32131–32143. doi: 10.1074/jbc.m114.594846.

24. Liu YH, Bu XL, Liang CR, Wang YR, Zhang T, Jiao SS, et al. An N-Terminal Antibody Promotes the Transformation of Amyloid Fibrils into Oligomers and Enhances the Neurotoxicity of Amyloid-Beta: The Dust-Raising Effect. Journal of neuroinflammation 2015, 12 (1). doi: 10.1186/s12974-015-0379-4.

25. pAb_2AT-L_, 6E10, and 4G8 all exhibited significant binding to Aβ_42_ by ELISA. 6E10 and 4G8 exhibited stronger binding than pAb_2AT-L_ (S5 Fig). Before treating the cells, the concentrations of all three antibodies were compared by absorbance at 280 nm using a nanodrop and confirmed to be in the same range, with pAb_2AT-L_ exhibiting a lower absorbance than 6E10 and 4G8.

26. Lin H, Dixon SG, Hu W, Hamlett ED, Jin J, Ergul A, Wang GY. P38 MAPK Is a Major Regulator of Amyloid Beta-Induced IL-6 Expression in Human Microglia. Molecular neurobiology 2022, 59 (9), 5284–5298. doi: 10.1007/s12035-022-02909-0.

27. Polini B, Ricardi C, Bertolini A, Carnicelli V, Rutigliano G, Saponaro F, et al. T1AM/TAAR1 System Reduces Inflammatory Response and β-Amyloid Toxicity in Human Microglial HMC3 Cell Line. International journal of molecular sciences 2023, 24 (14), 11569–11569. doi: 10.3390/ijms241411569.

28. Wei PC, Lee-Chen GJ, Chen CM, Chen Y, Lo YS, Chang KH. Isorhamnetin Attenuated the Release of Interleukin-6 from β-Amyloid-Activated Microglia and Mitigated Interleukin-6-Mediated Neurotoxicity. Oxidative Medicine and Cellular Longevity 2022, 2022, 3652402. doi: 10.1155/2022/3652402.

29. Wang WY, Tan MS, Yu JT, Tan L. Role of Pro-Inflammatory Cytokines Released from Microglia in Alzheimer’s Disease. Annals of Translational Medicine 2015, 3 (10), 7. doi: 10.3978/j.issn.2305-5839.2015.03.49.

30. Kayed R. Common Structure of Soluble Amyloid Oligomers Implies Common Mechanism of Pathogenesis. Science 2003, 300 (5618), 486–489. doi: 10.1126/science.1079469.

31. Kayed R, Head E, Sarsoza F, Saing T, Cotman CW, Necula M, et al. Fibril Specific, Conformation Dependent Antibodies Recognize a Generic Epitope Common to Amyloid Fibrils and Fibrillar Oligomers That Is Absent in Prefibrillar Oligomers. Molecular Neurodegeneration 2007, 2 (1), 18. doi: 10.1186/1750-1326-2-18.

32. Bigi A, Loffredo G, Cascella R, Cecchi C. Targeting Pathological Amyloid Aggregates with Conformation-Sensitive Antibodies. Current Alzheimer research 2020, 17 (8), 722–734. doi: 10.2174/1567205017666201109093848.

33. Bigi A, Napolitano L, Vadukul DM, Chiti F, Cecchi C, Aprile FA, Cascella RA Single-Domain Antibody Detects and Neutralises Toxic Aβ42 Oligomers in the Alzheimer’s Disease CSF. Alzheimer’s research & therapy 2024, 16 (1). doi: 10.1186/s13195-023-01361-z.

34. Kasturirangan S, Li L, Emadi S, Boddapati S, Schulz P, Sierks MR. Nanobody Specific for Oligomeric Beta-Amyloid Stabilizes Nontoxic Form. Neurobiology of Aging 2012, 33 (7), 1320–1328. doi: 10.1016/j.neurobiolaging.2010.09.020.

35. Meli G, Visintin M, Cannistraci I, Cattaneo A. Direct in Vivo Intracellular Selection of Conformation-Sensitive Antibody Domains Targeting Alzheimer’s Amyloid-β Oligomers. Journal of Molecular Biology 2009, 387 (3), 584–606. doi: 10.1016/j.jmb.2009.01.061.

36. Song C, Li H, Zheng C, Zhang T, Zhang Y. Dual Efficacy of a Catalytic Anti-Oligomeric Aβ42 ScFv Antibody in Clearing Aβ42 Aggregates and Reducing ABβBurden in the Brains of Alzheimer’s Disease Mice. Molecular neurobiology 2023, 60 (10), 5515–5532. doi: 10.1007/s12035-023-03406-8.

37. Zameer A, Kasturirangan S, Emadi S, Nimmagadda SV, Sierks MR. Anti-Oligomeric Aβ Single-Chain Variable Domain Antibody Blocks Aβ-Induced Toxicity against Human Neuroblastoma Cells. Journal of Molecular Biology 2008, 384 (4), 917–928. doi: 10.1016/j.jmb.2008.09.068.

38. Zhang Y, Chen X, Liu J, Zhang Y. The Protective Effects and Underlying Mechanism of an Anti-Oligomeric Aβ42 Single-Chain Variable Fragment Antibody. Neuropharmacology 2015, 99, 387–395. doi: 10.1016/j.neuropharm.2015.07.038.

39. Zhang X, Huai Y, Cai J, Song C, Zhang Y. Novel Antibody against Oligomeric Amyloid-β: Insight into Factors for Effectively Reducing the Aggregation and Cytotoxicity of Amyloid-Bβ Aggregates. International immunopharmacology 2019, 67, 176–185. doi: 10.1016/j.intimp.2018.12.014.

40. Lord A, Gumucio A, Englund H, Sehlin D, Sundquist VS, Söderberg L, et al. An Amyloid-β Protofibril-Selective Antibody Prevents Amyloid Formation in a Mouse Model of Alzheimer’s Disease. Neurobiology of Disease 2009, 36 (3), 425–434. doi: 10.1016/j.nbd.2009.08.007.

41. Hillen H, Barghorn S, Striebinger A, Labkovsky B, Müller R, Nimmrich V, et al. Generation and Therapeutic Efficacy of Highly Oligomer-Specific β-Amyloid Antibodies. The Journal of Neuroscience 2010, 30 (31), 10369–10379. doi: 10.1523/JNEUROSCI.5721-09.2010.

42. Several anti-Abeta antibodies or antibody fragments do exhibit protective effects at 1:1 or greater antibody-to-Abeta molar ratios.

43. Bodani RU, Sengupta U, Castillo-Carranza DL, Guerrero-Munoz MJ, Gerson JE, Rudra J, Kayed R. Antibody against Small Aggregated Peptide Specifically Recognizes Toxic Aβ-42 Oligomers in Alzheimer’s Disease. ACS chemical neuroscience 2015, 6 (12), 1981–1989. doi: 10.1021/acschemneuro.5b00231.

44. Gibbs E, Silverman JM, Zhao B, Peng X, Wang J, Wellington CL, et al. A Rationally Designed Humanized Antibody Selective for Amyloid Beta Oligomers in Alzheimer’s Disease. Scientific Reports 2019, 9 (1). doi: 10.1038/s41598-019-46306-5.

45. Lee EB. Targeting Amyloid-Beta Peptide (Abeta) Oligomers by Passive Immunization with a Conformation-Selective Monoclonal Antibody Improves Learning and Memory in Abeta Precursor Protein (APP) Transgenic Mice. Journal of Biological Chemistry 2006, 281 (7), 4292–4299. doi: 10.1074/jbc.m511018200.

46. Limbocker R, Mannini B, Cataldi R, Chhangur S, Wright AK, Kreiser RP, et al. Rationally Designed Antibodies as Research Tools to Study the Structure-Toxicity Relationship of Amyloid-β Oligomers. International Journal of Molecular Sciences 2020, 21 (12), 4542. doi: 10.3390/ijms21124542.

47. Sebollela A, Cline EN, Popova I, Luo K, Sun X, Ahn J, et al. A Human ScFv Antibody That Targets and Neutralizes High Molecular Weight Pathogenic Amyloid-Bβ Oligomers. Journal of Neurochemistry 2017, 142 (6), 934–947. doi: 10.1111/jnc.14118.

48. Stern AM, Liu L, Jin S, Liu W, Meunier AL, Ericsson M, et al. A Calcium-Sensitive Antibody Isolates Soluble Amyloid-β Aggregates and Fibrils from Alzheimer’s Disease Brain. Brain 2022. doi: 10.1093/brain/awac023.

49. Wacker J, Rönicke R, Westermann M, Wulff M, Reymann KG, Dobson CM, et al. Oligomer-Targeting with a Conformational Antibody Fragment Promotes Toxicity in Aβ-Expressing Flies. Acta neuropathologica communications 2014, 2 (1). doi: 10.1186/2051-5960-2-43.

50. It should be noted that pAb_2AT-L_ and other anti-Abeta antibodies have two antigen binding sites which may allow each antibody to bind two Abeta aggregates at once.

51. Abelein A, Abrahams JA, Danielsson J, Gräslund A, Jarvet J, Luo J, et al. The Hairpin Conformation of the Amyloid β Peptide Is an Important Structural Motif along the Aggregation Pathway. Journal of Biological Inorganic Chemistry 2014, 19 (4-5), 623-634. doi: 10.1007/s00775-014-1131-8.

52. Ciudad S, Puig E, Botzanowski T, Meigooni M, Arango AS, Do J, et al. Aβ(1-42) Tetramer and Octamer Structures Reveal Edge Conductivity Pores as a Mechanism for Membrane Damage. Nature Communications 2020, 11 (1), 3014. doi: 10.1038/s41467-020-16566-1.

53. Khaled M, Rönnbäck I, Ilag LL, Gräslund A, Strodel B, Österlund N. A Hairpin Motif in the Amyloid-β Peptide Is Important for Formation of Disease-Related Oligomers. Journal of the American Chemical Society 2023, 145 (33), 18340–18354. doi: 10.1021/jacs.3c03980.

54. Kreutzer AG, Guaglianone G, Yoo S, Parrocha CM, Ruttenberg SM, Malonis RJ, et al. Probing Differences among Aβ Oligomers with Two Triangular Trimers Derived from Aβ. 2023, 120 (22). doi: 10.1073/pnas.2219216120.

55. Sandberg A, Luheshi LM, Söllvander S, de Barros PT, Macao B, Knowles TPJ, et al. Stabilization of Neurotoxic Alzheimer Amyloid-β Oligomers by Protein Engineering. Proceedings of the National Academy of Sciences 2010, 107 (35), 15595–15600. doi: 10.1073/pnas.1001740107.

56. Sun Y, Kakinen A, Wan X, Moriarty N, Hunt CPJ, Li Y, et al. Spontaneous Formation of β-Sheet Nano-Barrels during the Early Aggregation of Alzheimer’s Amyloid Beta. Nano Today 2021, 38, 101125. doi: 10.1016/j.nantod.2021.101125.

57. Yu L, Edalji R, Harlan JE, Holzman TF, Lopez AP, Labkovsky B, et al. Structural Characterization of a Soluble Amyloid β-Peptide Oligomer. Biochemistry 2009, 48 (9), 1870–1877.

