## Supporting_information for "Antibodies raised against a structurally defined Aβ oligomer mimic protect human iPSC neurons from Aβ toxicity at sub-stoichiometric concentrations"

#### Table of Contents

**Supporting Figures** **S2**
- Figure S1. Rabbit antibodies and $A\beta_{42}$ in i[3]Neurons S2
- Figure S2. HMC3 microglia cells S2
- Figure S3. rabbit IgG TEM images S3
- Figure S4. Indirect ELISA of $pAb_{2AT-L}$, 6E10, and 4G8 against $A\beta_{42}$ S3
- Figure S5. Zoe images of i[3]neurons S4
- Table S1. Raw values and statistical analyses of $A\beta_{42}$ cell assays S4
- Table S2. Raw values and statistical analyses of $A\beta_{42}$ and antibody cell assays S5

**Materials and Methods** **S9**
- General Information S9
- Design of $pAb_{2AT-L}$ S9
- Generation of $pAb_{2AT-L}$ S9
- Affinity Purification of $pAb_{2AT-L}$ S10
- Preparation of $A\beta_{42}$ S11
- Indirect ELISAs S11
- Cell-Based Assays S12
- Thioflavin T Studies S14
- Transmission Electron Microscopy S15

**References and Notes** **S16**

#### Supporting Figures

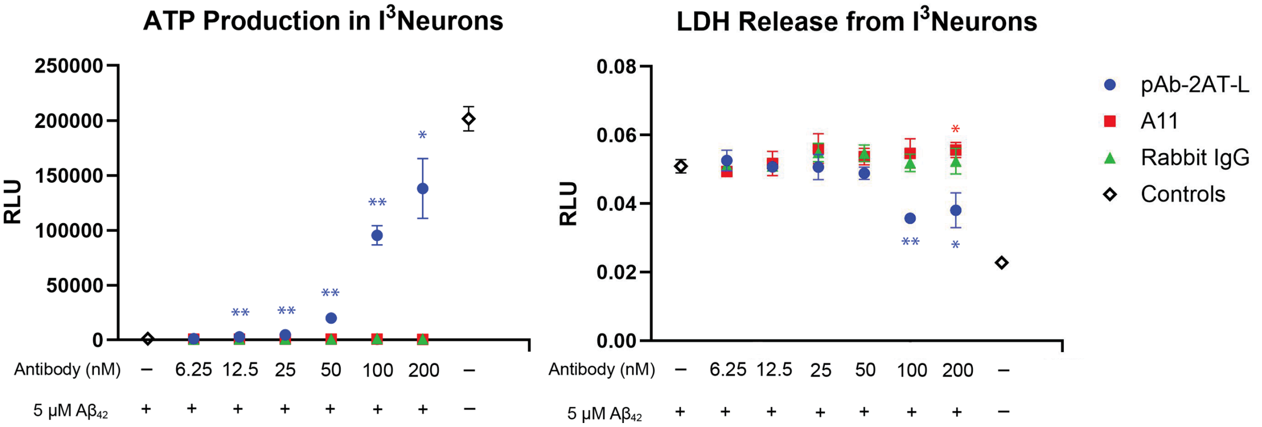


**Figure S1. Rabbit antibodies and Aβ42 in i³Neurons.** Graph of ATP production (left) and LDH release (right) by i³Neurons in the presence of 5 μM Aβ42 and varying concentrations of $pAb_{2AT-L}$, A11 (Fisher catalog #AHB0052), or a generic rabbit IgG (n=3 technical replicates). ** Significantly different ($p < 0.01$) from i³neurons treated with 5 μM Aβ42. *Significantly different ($p < 0.05$) from i³neurons treated with 5 μM Aβ42.

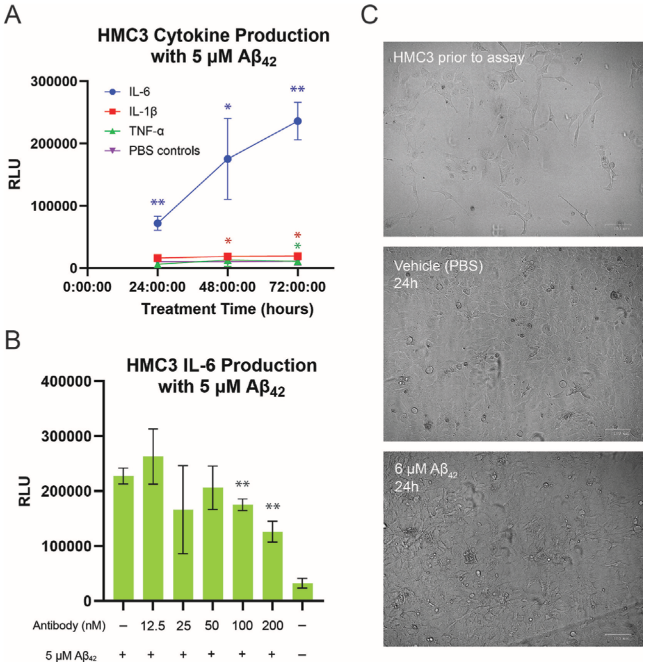


**Figure S2. HMC3 Microglia** (A) Graph of IL-6 production by HMC3 microglia cells in the presence of 5 μM Aβ42 and varying concentrations of $pAb_{2AT-L}$ as measured by the Promega™ Lumit™ IL-6 Human Immunoassay on the Promega™ GloMax® Discover plate reader (n=3 technical replicates). ** Significantly different ($p < 0.01$) from i³neurons treated with 5 μM Aβ42. (B) Graph of IL-6, IL-1β, and TNFα production by HMC3 microglia cells at different time points in the presence of 5 μM Aβ42 as measured by the Promega Lumit™ IL-6, Lumit™ IL-1β, and

Lumit™ TNF-α human immunoassays (n=4 technical replicates). ** Significantly different ($p <$ 0.01) from i[3]neurons treated with PBS (vehicle for $A\beta_{42}$). *Significantly different ($p < 0.05$) from i[3]neurons treated with PBS (vehicle for $A\beta_{42}$). (C) Images of HMC3 cells in culture prior to assaying (top), after 24h treatment with PBS (middle), and 6 μM $A\beta_{42}$ (bottom).

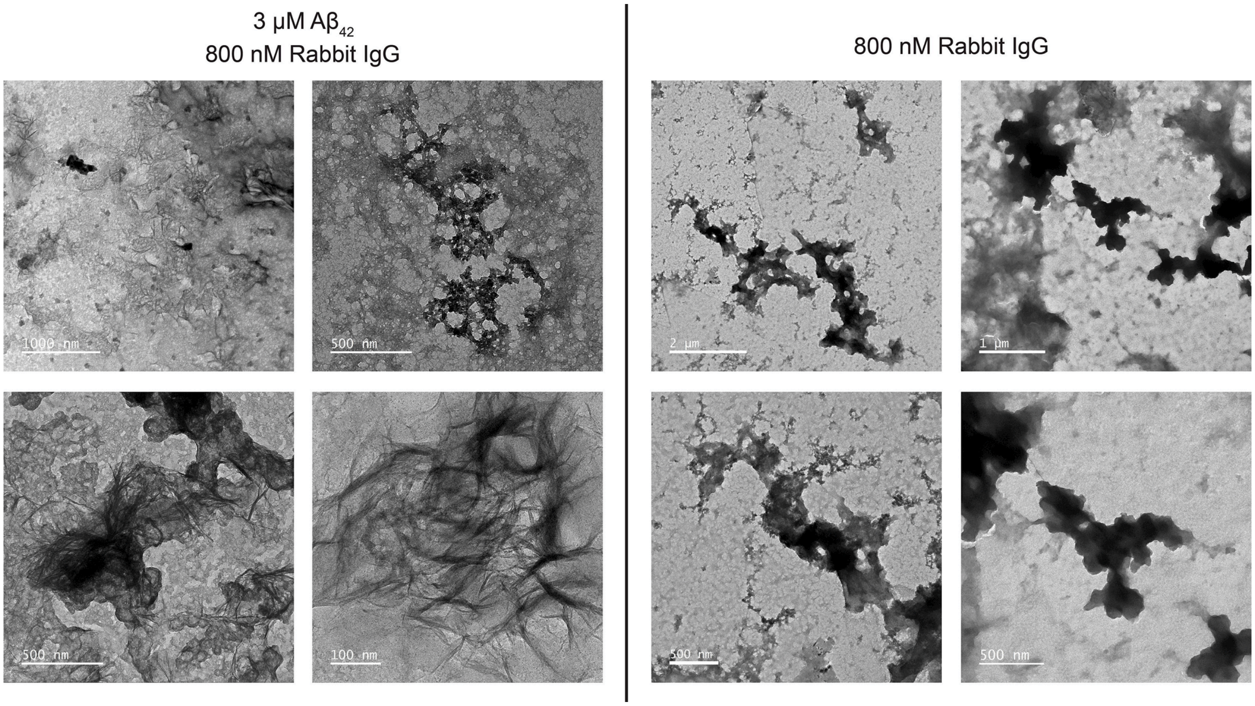


**Figure S3. TEM images with rabbit IgG.** Representative TEM images of $A\beta_{42}$ and a generic rabbit IgG (left) and generic rabbit IgG alone (right). Samples were all prepared in PBS and incubated for four hours prior to grid application. Samples applied to images and subsequently imaged as described.

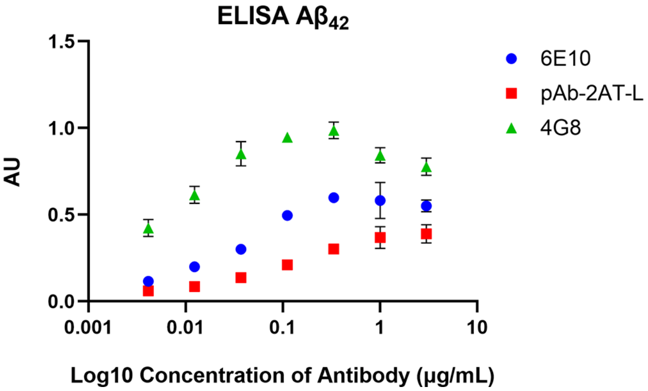


**Figure S4. Indirect ELISA of $pAb_{2AT-L}$, 6E10, and 4G8 against $A\beta_{42}$.** Graph depicting the binding of $pAb_{2AT-L}$, 6E10, and 4G8 to $A\beta_{42}$. Each well of a 96-well plate was treated with a 1 μM solution of $A\beta_{42}$ to coat the wells with $A\beta_{42}$. A 3-fold dilution series of $pAb_{2AT-L}$, 6E10, or 4G8 was then applied to the wells in triplicate, followed by an appropriate HRP-conjugated secondary antibody.

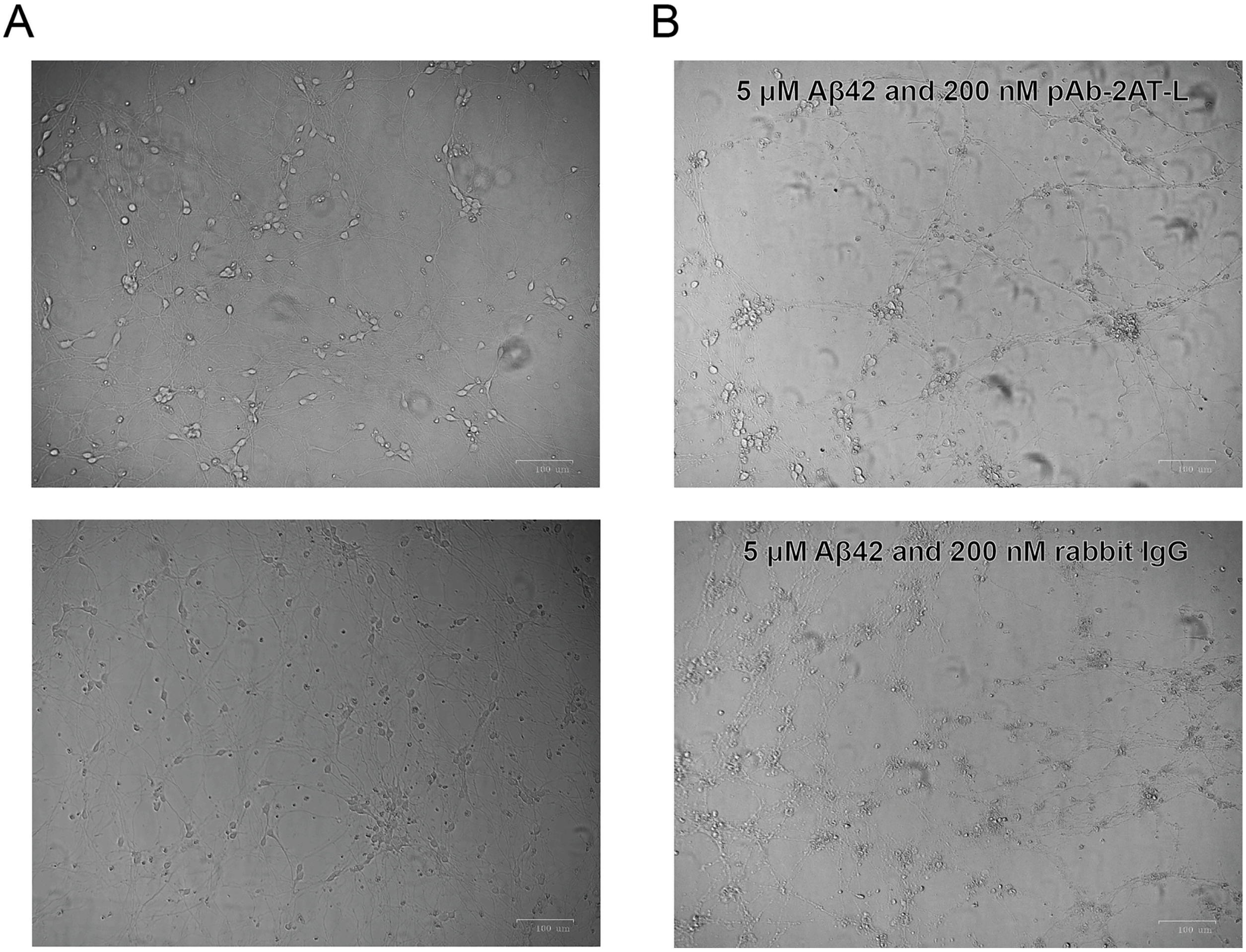


**Figure S5. ZOE images of i[3]Neurons** (A) treated with PBS (vehicle for $A\beta_{42}$) or (B) treated with 5 µM $A\beta_{42}$ and 200 nM antibody — $pAb_{2AT-L}$ (top) or generic rabbit IgG (bottom).

**Table S1. Table of raw values, averages, and t-tests from cell assays treating i[3]neurons with $A\beta_{42}$.**

| **Assay** | **Amount of $A\beta_{42}$** | **Rep 1** | **Rep 2** | **Rep 3** | **Rep 4** | **Rep 5** | **Rep 6** | **Avg** | **2-sample unequal variance t-test (*p*)** |
|---|---|---|---|---|---|---|---|---|---|
| ATP neurons (RLU) | 25 µM | 250 | 320 | 490 | 280 | 330 | 280 | 325 | 0.000242 |
| | 12.5 µM | 580 | 1030 | 1310 | 830 | 690 | 470 | 818.3333 | 0.000231 |
| | 6.25 µM | 4350 | 5300 | 4450 | 6150 | 5940 | 3990 | 5030 | 0.000159 |

| | | | | | | | | | |
|---|---|---|---|---|---|---|---|---|---|
| | 3.125 µM | 21300 | 20100 | 28000 | 28900 | 23700 | 16000 | 23000 | 2.89876E-08 |
| | 1.560 µM | 54400 | 46100 | 55300 | 65500 | 67300 | 56100 | 57450 | 7.00066E-07 |
| | 0.780 µM | 96500 | 97800 | 103000 | 106000 | 103000 | 101000 | 101216.7 | 0.000925 |
| | 0.390 µM | 85800 | 126000 | 112000 | 110000 | 104000 | 102000 | 106633.3 | 0.067834 |
| | 0.195 µM | 119000 | 117000 | 118000 | 127000 | 108000 | 107000 | 116000 | 0.382152 |
| | 0.098 µM | 119000 | 125000 | 109000 | 114000 | 109000 | 115000 | 115166.7 | 0.224514 |
| | 0 | 123000 | 117000 | 118000 | N/A | N/A | N/A | 119333.3 | Compared to all samples |
| LDH neurons (AU) | **25** µM | 0.0113 | 0.0143 | 0.011 | 0.0112 | 0.0115 | 0.0104 | 0.011617 | 0.000229 |
| | **12.5** µM | 0.0104 | 0.0102 | 0.0109 | 0.0114 | 0.0097 | 0.0101 | 0.01045 | 0.003988 |
| | **6.25** µM | 0.0109 | 0.0095 | 0.0097 | 0.0105 | 0.009 | 0.0069 | 0.009417 | 0.002349 |
| | **3.125** µM | 0.0095 | 0.007 | 0.0088 | 0.008 | 0.0092 | 0.0059 | 0.008067 | 0.01592625 |
| | **1.560** µM | 0.0066 | 0.0078 | 0.0041 | 0.0087 | 0.0082 | 0.0065 | 0.006983 | 0.124874776 |
| | 0.780 µM | 0.006 | 0.0067 | 0.0078 | 0.0073 | 0.0082 | 0.006 | 0.007 | 0.07843 |
| | 0.390 µM | 0.0075 | 0.0074 | 0.006 | 0.0056 | 0.0098 | 0.0064 | 0.007117 | 0.087877 |
| | **0.195** µM | 0.0036 | 0.0053 | 0.005 | 0.0052 | 0.0045 | 0.0039 | 0.004583 | 0.237041 |
| | **0.098** µM | 0.0058 | 0.0048 | 0.0065 | 0.0068 | 0.0067 | 0.0066 | 0.0062 | 0.313305 |
| | **0** | 0.0044 | 0.0061 | 0.0059 | | | | 0.005467 | Compared to all samples |

**Table S2. Table of raw values, averages, and t-tests from cell assays treating i[3]neurons and HMC3 microglia with A$\beta_{42}$ and anti- A$\beta$ antibodies.**

| **Assay** | **Concentration** | **Replicate 1** | **Replicate 2** | **Replicate 3** | **Avg** | **2-sample unequal variance t-test (*p*)** |
|---|---|---|---|---|---|---|
| ATP Neurons | Vehicle (PBS) | 215000 | 221000 | 201000 | 212333.3 | N/A |
| ATP Neurons 5 µM A$\beta_{42}$ (RLU) | Only A$\beta_{42}$ | 1200 | 840 | 950 | 996.6667 | Compared to all samples |
| | 200 nM pAb-2AT-L | 215000 | 219000 | 178000 | 204000 | 0.004105792 |
| | 100 nM pAb-2AT-L | 175000 | 176000 | 169000 | 173333.3 | 0.000155627 |

| | | | | | | |
|---|---|---|---|---|---|---|
| | 50 nM pAb-2AT-L | 83200 | 71800 | 83500 | 79500 | 0.002382263 |
| | 25 nM pAb-2AT-L | 15200 | 24600 | 22700 | 20833.33 | 0.020172524 |
| | 12.5 nM pAb-2AT-L | 4240 | 4900 | 7740 | 5626.667 | 0.048539044 |
| | 6.25 nM pAb-2AT-L | 1830 | 1580 | 1410 | 1606.667 | 0.020305871 |
| | 200 nM 6E10 | 1270 | 1350 | 1580 | 1400 | 0.047229895 |
| | 100 nM 6E10 | 1000 | 1150 | 1390 | 1180 | 0.304545989 |
| | 50 nM 6E10 | 990 | 960 | 1370 | 1106.667 | 0.553383451 |
| | 25 nM 6E10 | 1090 | 1040 | 1080 | 1070 | 0.563429704 |
| | 12.5 nM 6E10 | 870 | 1010 | 1070 | 983.3333 | 0.919529553 |
| | 6.25 nM 6E10 | 860 | 1050 | 820 | 910 | 0.540558296 |
| | 200 nM 4G8 | 5150 | 1770 | 2710 | 3210 | 0.157685392 |
| | 100 nM 4G8 | 2610 | 1430 | 1490 | 1843.333 | 0.150218419 |
| | 50 nM 4G8 | 1250 | 1130 | 1530 | 1303.333 | 0.127400896 |
| | 25 nM 4G8 | 1230 | 1060 | 1170 | 1153.333 | 0.279687515 |
| | 12.5 nM 4G8 | 1160 | 900 | 750 | 936.6667 | 0.727389197 |
| | 6.25 nM 4G8 | 650 | 580 | 830 | 686.6667 | 0.083169185 |
| LDH Neurons | Vehicle (PBS) | 0.0166 | 0.0156 | 0.0154 | 0.015867 | N/A |
| LDH Neurons 5 μM A$\beta_{42}$ (AU) | Only A$\beta_{42}$ | 0.0353 | 0.035 | 0.0366 | 0.035633 | Compared to all samples |
| | 200 nM pAb-2AT-L | 0.0192 | 0.0184 | 0.0232 | 0.020267 | 0.005138419 |
| | 100 nM pAb-2AT-L | 0.0171 | 0.0217 | 0.0237 | 0.020833 | 0.012891543 |
| | 50 nM pAb-2AT-L | 0.0278 | 0.0257 | 0.0244 | 0.025967 | 0.003463059 |
| | 25 nM pAb-2AT-L | 0.0335 | 0.0334 | 0.0345 | 0.0338 | 0.043844635 |
| | 12.5 nM pAb-2AT-L | 0.0319 | 0.0352 | 0.0379 | 0.035 | 0.754804362 |
| | 6.25 nM pAb-2AT-L | 0.0302 | 0.025 | 0.0269 | 0.027367 | 0.023435481 |
| | 200 nM 6E10 | 0.037 | 0.0342 | 0.0366 | 0.035933 | 0.783485076 |
| | 100 nM 6E10 | 0.037 | 0.0355 | 0.0363 | 0.036267 | 0.38905776 |
| | 50 nM 6E10 | 0.0368 | 0.035 | 0.0354 | 0.035733 | 0.898295002 |

| | | | | | | |
|---|---|---|---|---|---|---|
| | 25 nM 6E10 | 0.035 | 0.0344 | 0.0363 | 0.035233 | 0.620430854 |
| | 12.5 nM 6E10 | 0.0348 | 0.0356 | 0.0342 | 0.034867 | 0.297197599 |
| | 6.25 nM 6E10 | 0.0305 | 0.0336 | 0.0294 | 0.031167 | 0.055975855 |
| | 200 nM 4G8 | 0.0424 | 0.0374 | 0.0343 | 0.038033 | 0.417048709 |
| | 100 nM 4G8 | 0.0378 | 0.0363 | 0.0364 | 0.036833 | 0.156829082 |
| | 50 nM 4G8 | 0.0387 | 0.0369 | 0.0364 | 0.037333 | 0.1253391 |
| | 25 nM 4G8 | 0.0369 | 0.0363 | 0.0362 | 0.036467 | 0.226412319 |
| | 12.5 nM 4G8 | 0.0394 | 0.0378 | 0.0368 | 0.038 | 0.068648713 |
| | 6.25 nM 4G8 | 0.0296 | 0.0259 | 0.0304 | 0.028633 | 0.026401684 |
| ATP Neurons | Vehicle (PBS) | 189000 | 210000 | 206000 | 201666.7 | N/A |
| ATP Neurons 5 μM A$\beta_{42}$ (RLU) | Only A$\beta_{42}$ | 1560 | 820 | 810 | 1063.333 | Compared to all samples |
| | 200 nM pAb-2AT-L | 149000 | 158000 | 107000 | 138000 | 0.012900905 |
| | 100 nM pAb-2AT-L | 85100 | 99600 | 101000 | 95233.33 | 0.002845161 |
| | 50 nM pAb-2AT-L | 18500 | 19100 | 22200 | 19933.33 | 0.002592076 |
| | 25 nM pAb-2AT-L | 4910 | 4490 | 4960 | 4786.667 | 0.000648166 |
| | 12.5 nM pAb-2AT-L | 2970 | 2950 | 2640 | 2853.333 | 0.009498503 |
| | 6.25 nM pAb-2AT-L | 1110 | 1180 | 980 | 1090 | 0.925449051 |
| | 200 nM A11 | 540 | 500 | 680 | 573.3333 | 0.182501208 |
| | 100 nM A11 | 520 | 590 | 880 | 663.3333 | 0.244954964 |
| | 50 nM A11 | 610 | 620 | 660 | 630 | 0.222770616 |
| | 25 nM A11 | 650 | 640 | 700 | 663.3333 | 0.248120178 |
| | 12.5 nM A11 | 810 | 870 | 740 | 806.6667 | 0.410284494 |
| | 6.25 nM A11 | 770 | 730 | 790 | 763.3333 | 0.350412919 |
| | 200 nM Rab IgG | 610 | 880 | 1110 | 866.6667 | 0.539779431 |
| | 100 nM Rab IgG | 1210 | 1180 | 1360 | 1250 | 0.533497189 |
| | 50 nM Rab IgG | 950 | 870 | 1200 | 1006.667 | 0.847592589 |
| | 25 nM Rab IgG | 800 | 960 | 1070 | 943.3333 | 0.683487863 |

| | | | | | | |
|---|---|---|---|---|---|---|
| | 12.5 nM Rab IgG | 830 | 1090 | 880 | 933.3333 | 0.660060836 |
| | 6.25 nM Rab IgG | 580 | 740 | 790 | 703.3333 | 0.281850838 |
| LDH Neurons | Vehicle (PBS) | 0.0228 | 0.0216 | 0.0239 | 0.022767 | N/A |
| LDH Neurons 5 μM Aβ42 (AU) | Only Aβ42 | 0.0529 | 0.049 | 0.0508 | 0.0509 | Compared to all samples |
| | 200 nM pAb-2AT-L | 0.0355 | 0.0346 | 0.0439 | 0.038 | 0.035735572 |
| | 100 nM pAb-2AT-L | 0.0356 | 0.0357 | 0.0357 | 0.035667 | 0.005395934 |
| | 50 nM pAb-2AT-L | 0.0509 | 0.0477 | 0.0478 | 0.0488 | 0.244853585 |
| | 25 nM pAb-2AT-L | 0.0475 | 0.0547 | 0.0497 | 0.050633 | 0.918797077 |
| | 12.5 nM pAb-2AT-L | 0.052 | 0.0507 | 0.0494 | 0.0507 | 0.890719776 |
| | 6.25 nM pAb-2AT-L | 0.0522 | 0.0497 | 0.0557 | 0.052533 | 0.481668002 |
| | 200 nM A11 | 0.0532 | 0.0563 | 0.0574 | 0.055633 | 0.049311436 |
| | 100 nM A11 | 0.0508 | 0.0537 | 0.0593 | 0.0546 | 0.275825661 |
| | 50 nM A11 | 0.0511 | 0.0539 | 0.056 | 0.053667 | 0.205142344 |
| | 25 nM A11 | 0.0519 | 0.0557 | 0.0605 | 0.056033 | 0.163778481 |
| | 12.5 nM A11 | 0.0531 | 0.0476 | 0.0543 | 0.051667 | 0.76509396 |
| | 6.25 nM A11 | 0.0507 | 0.0485 | 0.0489 | 0.049367 | 0.32124822 |
| | 200 nM Rab IgG | 0.055 | 0.054 | 0.0481 | 0.052367 | 0.588445681 |
| | 100 nM Rab IgG | 0.049 | 0.0533 | 0.0534 | 0.0519 | 0.616715288 |
| | 50 nM Rab IgG | 0.0573 | 0.0526 | 0.0541 | 0.054667 | 0.105535833 |
| | 25 nM Rab IgG | 0.0524 | 0.0578 | 0.0544 | 0.054867 | 0.11734549 |
| | 12.5 nM Rab IgG | 0.0504 | 0.0504 | 0.0522 | 0.051 | 0.94242587 |
| | 6.25 nM Rab IgG | 0.0507 | 0.0524 | 0.0504 | 0.051167 | 0.848718665 |
| IL-6 HMC3 | Vehicle (PBS) | 26300 | 39000 | N/A | 32650 | N/A |
| IL-6 HMC3 5 μM Aβ42 (RLU) | Only Aβ42 | 211000 | 237000 | 235000 | 227666.7 | Compared to all samples |
| | 200 nM pAb-2AT-L | 147000 | 110000 | 122000 | 126333.3 | 0.002311801 |
| | 100 nM pAb-2AT-L | 175000 | 186000 | 165000 | 175333.3 | 0.009067724 |
| | 50 nM pAb-2AT-L | 181000 | 252000 | 186000 | 206333.3 | 0.456421006 |

| | | | | | | |
|---|---|---|---|---|---|---|
| | 25 nM pAb-2AT-L | 212000 | 73800 | 213000 | 166266.7 | 0.314485693 |
| | 12.5 nM pAb-2AT-L | 237000 | 231000 | 321000 | 263000 | 0.34814919 |

### Materials and Methods

#### General Information

All chemicals were used as received unless otherwise noted. Deionized water (18 MΩ) was obtained from a Barnstead NANOpure Diamond water purification system. 6E10 and 4G8 were purchased from Biolegend (cataolog #803016 and #800702, respectively). The generic IgG rabbit antibody was purchased from ThermoFisher Scientific (catalog #31235).

#### Design of $pAb_{2AT\text{-}L}$

2AT-L consists of three cyclic β-hairpin peptides covalently linked by three disulfide bonds to form a triangular trimer. The individual peptides that form this trimer were designed based on crystal structures of peptides that mimic the 17–36 Aβ β-hairpin originally reported by Hard and coworkers in 2007 [1]. These peptides all have a propensity to assemble into trimers which further assemble into higher-order oligomers [2-5]. The monomers that comprise 2AT-L, are each composed of Aβ residues 17 through 36 connected at the C and N termini by a δ-linked ornithine turn unit. This linkage conformationally constrains the peptide in a β-hairpin conformation. Each peptide also contains an N-methyl on the backbone of the peptide at Phe20 to prevent fibrillization, a mutation at Met35 to α-linked ornithine to aid in structural elucidation, and two cysteine mutations at positions 17 and 21 to allow for disulfide bond formation. Upon oxidation, this peptide readily forms triangular trimers (2AT-L) [6].

The structure of 2AT-L was elucidated at atomic resolution by X-ray crystallography, and further characterized by SDS-PAGE, size-exclusion chromatography, and dynamic light scattering, revealing a propensity to form higher-order assemblies in solution [6]. Cell assays showed that 2AT-L elicits LDH release, decreases ATP production, and activates caspase-3/7 mediated apoptosis in SH-SY5Y neuroblastoma cells [6]. We have also examined the effects of 2AT-L on iPSC-derived neurons and have observed decreased ATP production and activation of caspase 3/7 activation at concentrations of 10 μM or greater.

#### Generation of $pAb_{2AT\text{-}L}$

$pAb_{2AT\text{-}L}$ was generated by Pacific Immunology (Ramona, CA, www.pacificimmunology.com) using standard custom antibody production procedures. Briefly, 2AT-L was conjugated to the carrier protein keyhole limpet hemocyanin (KLH) using standard EDC conjugation chemistry. The EDC was obtained from Thermo Scientific (catalog # 22980) and the conjugation was performed according to manufacturer's instructions in MES buffer at an acidic pH of 5. We have previously found that at acidic pH, a related covalently stabilized trimer does not assemble to form higher-order assemblies [4]. It is thus likely that 2AT-L does not assemble to form higher-order structures under the conditions of the conjugation reaction to KLH. Two New Zealand white rabbits, 9–10 weeks of age, were used to generate $pAb_{2AT\text{-}L}$. Rabbits were primed with 200 µg of the 2AT-L-KLH conjugate in AdjuLite Complete Freund's Adjuvant (CFA) (Cat. #A5001), by subcutaneous injection using a 19-gauge needle in the neck/shoulder region. On days 21, 42, and 70 the rabbits received booster injections of 100 µg of the 2AT-L-KLH conjugate in AdjuLite Incomplete Freund's Adjuvant (IFA) (Cat. #A5002). Blood was drawn from the central ear artery using a 19-gauge needle on days 0, 49, 63, 77, and 91.

As whole blood was collected, it was diluted in Anticoagulant Citrate Dextrose Solution USP (ACD) Solution A (0.80 g citric acid monohydrate, 2.45 g dextrose monohydrate, and 2.20 g of sodium citrate dihydrate in 100 mL) in a 1:10 ratio. The blood-ACD solution was diluted 1:1 with sterile PBS containing 2.5 mM EDTA and 0.5% BSA. 20 mL of StemCell Lymphoprep was added to a 50-mL StemCell SepMate-50 separation tube, then 30 mL of the blood and ACD solution was added over the Lymphoprep, then centrifuged for 30 minutes at 3,000 RPM. The supernatant (plasma) was retrieved and decanted once the PBMCs have been spun down to form a pellet. The plasma was then used in the affinity purification as described below.

#### Affinity purification of $pAb_{2AT\text{-}L}$

$pAb_{2AT\text{-}L}$ was purified from the plasma using affinity chromatography. To create the affinity chromatography column, 10 mg of the 2AT-L TFA salt was dissolved in 2 mL DMSO. The 2AT-L/DMSO solution was then added to 10 mL PBS (pH 7.4) to create a 1 mg/mL 2AT-L solution in PBS with 20% DMSO. The 1 mg/mL 2AT-L solution was then added to 750 mg dry NHS-activated agarose resin (Thermo Scientific, catalog # 26197) in a 15 mL polypropylene conical tube. The 2AT-L/agarose suspension was then continuously inverted using a tube rotator for two hours at room temperature. After the two-hour incubation, the agarose was transferred to a Bio-Rad Poly-Prep chromatography column (10 mL) and washed 3x with PBS. The agarose was then incubated in 5 mL 1 M Tris buffer (pH 8.5) for 1 hour at room temperature on a tube rotator to cap unreacted NHS-ester groups, and then washed 3x with PBS.

For the affinity purification of $pAb_{2AT\text{-}L}$ from the plasma, the 2AT-L-agarose was transferred to a 50 mL CrystalCruz® glass chromatography column. The plasmas from a single production bleed from both rabbits were combined and then filtered through 0.45 µm syringe filters. The filtered plasma was then added to the column containing the 2AT-L-agarose and rocked on a rocker for ~3 hours at room temperature or overnight at 4 ºC. Next, the plasma was drained from the column into a 50 mL polypropylene conical tube, and the 2AT-L-agarose was washed with ice-cold PBS. Washing was performed with 50 mL portions of ice-cold PBS until the absorbance at 214 nm of the wash buffer eluent measured less than 0.05 on a NanoDrop One/One$^{c}$ Microvolume UV-Vis Spectrophotometer.

$pAb_{2AT\text{-}L}$ was then eluted from the 2AT-L-agarose using ice-cold 0.2 M glycine buffer (pH 1.85). The $pAb_{2AT\text{-}L}$ elution was performed by adding the glycine buffer in 1 mL portions and then collecting the eluent into 1.7 mL microcentrifuge tubes containing 0.5 mL 1 M Tris buffer (pH 8.5). Typically, 10–15 1-mL portions of the glycine buffer were sufficient to elute all of $pAb_{2AT\text{-}L}$ from the

2AT-L-agarose. The eluents were then combined and transferred to a 30 kDa molecular weight cutoff filter (30K MWCOF) (Thermo Scientific catalog # 88531) and buffer exchange with PBS was performed. The 30K MWCOF was centrifuged in a swinging bucket centrifuge at 3800 x g until the volume fell below 1 mL, after which the buffer was replenished with ice-cold PBS up to the 20 mL marker on the 30K MWCOF. This process was repeated at least six times. For the final centrifugation step, the $pAb_{2AT-L}$ volume was allowed to reach ~0.5 mL in the 30K MWCOF and was then transferred to a 1.7 mL microcentrifuge tube. The $pAb_{2AT-L}$ concentration was determined using a BCA assay and then adjusted to 1 mg/mL with PBS. Typical affinity purifications yielded 1–3 mg of $pAb_{2AT-L}$ from 50 mL of antisera. The affinity purified $pAb_{2AT-L}$ was then portioned into 50 µL aliquots, which were stored at -80 ºC and thawed as needed. Once thawed, the aliquot was stored at 4 ºC.

#### Preparation of $A\beta_{42}$

A 1 mg portion of recombinantly expressed $A\beta_{42}$ as the ammonium salt was purchased from rPeptide (catalog# A-1167-2) and received as a fluffy lyophilized solid in a glass amber vial. The 1 mg $A\beta_{42}$ portion was dissolved with 1 mL of 2 mM NaOH to create a 1 mg/mL $A\beta_{42}$ solution. The 1 mg/mL $A\beta_{42}$ solution was then sonicated in a water bath sonicator for 5 minutes. After sonication, 0.02 µmol aliquots of $A\beta_{42}$ were prepared by transferring 92.6 µL portions of the 1 mg/mL $A\beta_{42}$ solution to low-binding microcentrifuge tubes (Axygen, catalog# MCT-175-L-C) containing a hole in the lid of the tube created by puncturing the lid with a 22-gauge needle. The aliquots were then frozen on dry ice for 1 hour, transferred to a lyophilization vessel, and lyophilized overnight. The next day, the aliquots were removed from the lyophilizer and each microcentrifuge tube was immediately transferred to its own 50 mL conical tube. The 50 mL conical tubes were sealed by tightening the lid and stored at -80 °C until use.

#### Indirect ELISAs

Indirect ELISA was used to verify the binding of $pAb_{2AT-L}$, 4G8, and 6E10 to $A\beta_{42}$. Three technical replicates were performed for each experiment. Each aspiration and washing step in the ELISA procedure was performed using a Fisherbrand™ accuWash™ Microplate Washer (catalog # 14-377-577).

**Coating the wells of the ELISA plates.** A 0.02 µmol aliquot of $A\beta_{42}$ (prepared as described above) was removed from the -80 °C freezer and allowed to equilibrate to room temperature. The aliquot was then dissolved in 92.6 µL of deionized water to create a 216 µM solution of $A\beta_{42}$. A 1 µM solution of $A\beta_{42}$ in carbonate buffer (15 mM $Na_2CO_3$, 35 mM $NaHCO_3$, 0.02% (w/v) sodium azide, pH 9.5) was then prepared from the 216 µM solution. The 1 µM $A\beta_{42}$ solution was then poured into a reagent reservoir and a multichannel pipette was used to transfer 50 µL to the appropriate wells of a Thermo Scientific™ Maxisorp 96-well plate (catalog# 12-565-135). The 96-well plate was then sealed with an adhesive 96-well plate seal (Axygen, catalog# PCR-SP) and incubated overnight (~16 hours) at room temperature on a rotating shaker set to 90 RPM.

**Treating the ELISA plates with $pAb_{2AT-L}$, 4G8, or 6E10.** The next day, 20 mL of 1% BSA was prepared by adding 200 mg of BSA (Fraction V) (Fisher BioReagents™, catalog # BP1600-100) to 20 mL PBS. The solution of $A\beta_{42}$ was aspirated from the wells of the 96-well plate and then washed 1x with PBST (10 mM $Na_2HPO_4$, 1.8 mM $KH_2PO_4$, 137 mM NaCl, 2.7 mM KCl, 0.5% Tween-20). A multichannel pipette was then used to transfer 75 µL of 1% BSA to each well, the plate was sealed, and incubated for at least 1 hour at room temperature on a rotating shaker set to 90 RPM to block uncoated sites in the wells. During the last 15 minutes of the blocking step, a 3 µg/mL $pAb_{2AT-L}$ solution, 3 µg/mL 6E10 solution, and 3 µg/mL 4G8 solution were prepared by diluting a 636 nL of a 1.18 mg/ml ($pAb_{2AT-L}$) or 750 nL 1.0 mg/ml (6E10 and 4G8) stock solution into 250 µL of 1% BSA in PBS. After the blocking step, the wells were aspirated and washed 3x with PBST. Using a multi-channel pipette, 50 µL of 1% BSA was then added to the wells in the bottom seven rows of the 96-well plate (rows B–H). 75 µL of the 3 µg/mL $pAb_{2AT-L}$, 6E10, and 4G8 solutions were added to their respective wells in the top row (row A). A multi-channel pipette was then used to create a three-fold dilution series of the antibodies by transferring 25 µL of the solutions from row A to row B and then mixing up and down 8 times. 25 µL was then transferred from row B to row C, and so on, until the last 25 µL had been transferred to row H. The 96-well plates were then sealed with an adhesive plate seal and incubated for 2 hours at room temperature on a rotating shaker set to 90 RPM.

**Treating the ELISA plate with the secondary antibodies.** After the 2-hour incubation, the primary antibody solutions were aspirated and the wells were washed 3x with PBST. A 50 µL portion of AffiniPure Goat Anti-Rabbit IgG (H+L) conjugated to horse radish peroxidase (GαR-HRP; Jackson ImmunoResearch, catalog # 111-035-144) diluted 1:10,000 in 1% BSA was then added to the wells that received $pAb_{2AT-L}$; a 50 µL portion of AffiniPure Goat Anti-Mouse IgG (H+L) conjugated to horse radish peroxidase (GαM-HRP; Jackson ImmunoResearch, catalog # 115-035-146) diluted 1:10,000 in 1% BSA was added to the wells that received 6E10 and 4G8. The 96-well plate was then sealed with an adhesive plate seal and incubated for 1 hour at room temperature on a rotating shaker set to 90 RPM.

**Developing the ELISA plates.** After the 1-hour incubation, the secondary antibody solutions were aspirated and the wells were washed 3x with PBST. A 50 µL portion of 3,3',5,5'-tetramethybenzidine (TMB) (Millipore Sigma, catalog # ES001-500ML) was then added to each well and allowed to react until the blue color reached a sufficient hue. A 50 µL portion of 1 M aqueous HCl was then added to each well to quench the reaction and the absorbance was measured at 450 nm using a MultiSkan GO plate reader. The absorbance readings for the three replicates were averaged and the standard deviations were calculated using GraphPad Prism. The data were then plotted and fit using GraphPad Prism to estimate the $EC_{50}$ values.

#### Cell-Based Assays

**Differentiation of NGN2 iPSCs into cortical neurons.** Ngn2 iPSCs were obtained from the UCI Sue and Bill Gross Stem Cell Research Center, courtesy of the Blurton-Jones laboratory and differentiated according to the protocols published by Gan and coworkers [7]. An aliquot of Ngn2 iPSCs were thawed in 10 mL pre-warmed mTeSR media treated with 10 µL of 10 mM rock inhibitor (final concentration 10 μM) and centrifuged at 200g for 4 minutes. The media was aspirated and replaced with fresh mTesR with rock inhibitor and plated at a density of about 500,000 cells per ml on a Matrigel coated 6-well plate (2 ml per well). Cells were incubated at 37 °C with 5% CO2 and cell media was aspirated and replaced daily with mTesR (without rock inhibitor) until the cell density reached 80% confluency. Cells were then passaged by aspirating the media, treating with accutase, centrifuging, and replating in fresh mTesR with rock inhibitor.

Once the cell density again reached 80 % confluency, cells were passaged and resuspended in neuronal predifferentiation media (Knockout DMEM:F12, 1x N2, 1x non-essential amino acids, 10 ng/ml BDNF, 10 ng/ml NT3, 1 ug/ml mouse laminin, 2 ug/ml doxyclyine) with rock inhibitor (10 uM), plated in a new Matrigel coated 6-well plate at the same density in 2ml of media per well, and incubated at 37 °C. The following two days media was aspirated and replaced. Cells were then either frozen in Knockout DMEM with 10% DMSO for later maturation or passaged and resuspended in neuronal maturation media (1:1 neurobasal A media and DMEM f:12, 1x B27 0.5x N2, 1x non-essential amino acids, 0.5x glutamax 10 ng/ml BDNF, 10 ng/ml NT3, 1 ug/ml mouse laminin, 2 ug/ml doxyclyine).

96-well black, clear bottom, tissue culture treated plates were coated in 0.5 mg/ml poly-D-lysine solution in Dubeccos Phosphate buffered saline. After one hour, the solution was removed and the plates were washed twice with DPBS and left to dry for two hours. Cells were plated for maturation on the inner 60 wells of the plate at a density of 200000 cells per ml. Media was added to the outer wells of the plate to saturate the environment. After one week of maturation, media was gently removed and replaced with maturation media containing the compounds of interest, but lacking doxycycline. The neurons were incubated for 72 hours before assaying.

**HMC3 microglia culture.** HMC3 microglia cells were purchased from ATCC (Cat# CRL-3304). Upon arrival, cells were stored at -80 °C for 2 days and then thawed in a bead bath at 37 °C. The cells were transferred to a 15 mL conical containing 9 mL of pre-warmed EMEM with 10 % fetal bovine serum. The conical was centrifuged at 125 g for 5 min, the supernatant was aspirated, and the cell pellet was redissolved in 1 mL of media and transferred to a 75 $cm^2$ cell culture flask containing 19 mL of media. The cells were incubated at 37 °C with 5% $CO_2$ until reaching 80 % confluence (three to five days) before passaging with trypsin to a new cell culture flask. On the subsequent passage, the cells were plated on the inner 60 wells of a black, clear-bottom, 96-well cell culture plate at a density of one hundred thousand cells per mL and incubated overnight before treatment. The cells were incubated for 72 hours before assaying.

**Preparation of reagents for cell assays.** Antibodies were diluted in PBS to 10X of the desired final concentration and spin filtered. In a replica 96-well plate antibodies were diluted in a series of two-fold dilutions with PBS in triplicate to get final volumes of 10 μL per well. The positive and negative controls contained just 10 μL of PBS. For cell assays, two aliquots of 0.02 μmoles of Aβ42 were dissolved in 7.2 mL of maturation media to obtain a concentration of 5.56 μM. 90 μL of 5.56 μM Aβ42 in maturation media was added to each well of the replica plate, except the negative control which had media without Aβ42. Upon receipt, CellTiter-Glo® 2.0 cell viability reagent as thawed, aliquoted, and stored at -80 °C. For the assay, one 4 μL aliquot was thawed to room temperature before use. The CyQuant™ LDH substrate and buffer were stored at -20 °C. For the assay, the LDH buffer was equilibrated to room temperature, dissolved in 11.4 ml of nanopure water, and vortexed. 600 μL of LDH substrate was equilibrated to room temperature and then added to the reconstituted LDH buffer and vortexed before use. The Anti-hIL-6 antibodies were stored at -20 °C upon receipt. Immediately before the assay, 12 μL of the SmBiT and 12 μL of the LgBiT antibodies were added to 5.976 mL of cell culture media to make 6 mL of the 2X Anti-hIL-6 antibody mixture. The Lumit detection reagent was made by adding 80 μL of Lumit Detection Substrate B to 1.52 mL of Lumit Detection Buffer B.

**CellTiter-Glo® 2.0 Cell Viability Assay and CyQuant™ LDH Cytotoxicity Assay.** Cell assays were performed according to manufacturers' instructions. After 72 hours of incubation, the cells were removed from the incubator and allowed to equilibrate to room temperature for 30 minutes. 50 μL of media per well was removed and transferred to a clear 96-well plate. 50 μL of LDH reagent was added to the clear plate, briefly mixed, and incubated in the dark for 1 hour. The absorbance of each well was measured at 490 nm on a Thermo Scientific MultiSkan GO plate reader. The remaining 50 μL of media containing the cells was treated with 50 μL per well of CellTiter-Glo® 2.0 viability reagent. The plate was incubated at room temperature with orbital shaking for 15 minutes at 80 RPM. The luminescence from each well was then measured on a Promega™ GloMax® Discover Microplate Reader. The absorbance and luminescence measurements for the three replicates of each treatment group were averaged and the standard deviations were calculated using GraphPad Prism for each assay. The data were then plotted using GraphPad Prism.

**Lumit™ Il-6 (Human) Immunoassay.** Cell assays were performed according to manufacturers' instructions. After 72 hours of incubation, the cells were treated with 200 μL per well of 2X Anti-hIL-6 antibody mixture and returned to the incubator for 45 minutes. The cells were then treated with 40 μL per well of 5X Lumit detection reagent, briefly mixed, and incubated at room temperature for 5 minutes before measuring luminescence with a Promega™ GloMax® Discover Microplate Reader. The data were then plotted using GraphPad Prism.

#### Thioflavin T assay on $A\beta_{42}$

The ThT assay was performed on 3 μM $A\beta_{42}$ in PBS at pH 7.4 (10 mM $Na_2HPO_4$, 1.8 mM $KH_2PO_4$, 137 mM NaCl, 2.7 mM KCl) containing 10 μM ThT in the presence of a dilutions series of $pAb_{2AT\text{-}L}$ (800–210 nM, 0nM). A 0.02 μmol aliquot of $A\beta_{42}$ was removed from the -80 °C freezer and allowed to equilibrate to room temperature. During this equilibration time, an 10 μM solution of thioflavin T (ThT) was prepared in PBS at pH 7.4 (10 mM $Na_2HPO_4$, 1.8 mM $KH_2PO_4$, 137 mM NaCl, 2.7 mM KCl) and a serial dilution series of 10x concentrations of $pAb_{2AT\text{-}L}$ were prepared in deionized water. The assay was performed in triplicate in Corning® 96-well Half Area Black/Clear Flat Bottom Polystyrene NBS Microplates (product# 3881) at 25 °C under quiescent conditions.

**Preparation of 10 μM ThT in PBS.** To prepare the 10 μM ThT solution in PBS, a concentrated solution of ThT (1–3 mM) was prepared in a 15 mL polypropylene conical tube by adding ~5–7 mg of ThT (TCI Chemicals, catalog# T0558) to 7–10 mL PBS. The concentrated ThT solution was sonicated in a water bath sonicator for ~5 minutes and then passed through a 0.2 μm nylon syringe filter into a new 15 mL conical tube. The concentration of the concentrated ThT solution was determined photospectrometrically by first preparing 2 mL of a 1:200 diluted ThT solution (0.010 mL of the concentrated ThT solution into 1.990 mL PBS). The absorbance of the diluted ThT solution was then measured at 412 nm in a 1 cm quartz cuvette and the concentration was calculated using an estimated extinction coefficient (ε) of 36,000 $M^{-1}$ $cm^{-1}$. The concentration of the diluted ThT solution was multiplied by 200 to calculate the concentration of the concentrated ThT solution. A 10 mL portion of 10 μM ThT in PBS was prepared by diluting an appropriate volume of the concentrated ThT solution with PBS. The 10 μM ThT solution was then kept on ice until used.

**Preparation of pAb2AT-L.** A 1.25-fold dilution series containing 10x concentrations of $pAb_{2AT-L}$ (8–2.1 μM, 0 μM) was prepared in row A of the 96-well plate. To prepare these dilution series, 30 μL of an 8 μM solution of $pAb_{2AT-L}$ in PBS was added to well A1, and 15 μL PBS was added to wells A2–A8. The serial dilution was then performed by transferring 15 μL from well A1 to well A2 and then mixing by pipetting up and down 8–12 times, and so on, stopping on well A7. Well A8 did not receive $pAb_{2AT-L}$ and only contained 15 μL of PBS. A 12-channel pipette was then used to transfer 4 μL portions of the $pAb_{2AT-L}$ solutions row A to rows B,C, and D. After the dilution series was prepared and aliquoted to the respective rows, the 96-well assay plate was kept on ice for the remainder of the experimental setup.

**Adding $A\beta_{42}$ to the ThT assay plate.** After the 10 μM ThT solution was prepared and the $pAb_{2AT-L}$ dilution series was completed, a 3.33 μM solution of $A\beta_{42}$ was prepared in 10 μM ThT. To prepare the 3.33 μM solution of $A\beta_{42}$, the equilibrated 0.02 μmol $A\beta_{42}$ aliquot was first dissolved in 60 μL deionized water to create a 333 μM solution of $A\beta_{42}$ and sonicated in a water bath sonicator for 5 minutes. During the sonication time, 5.94 mL of ice-cold 10 μM ThT was transferred to a 15 mL conical tube and kept on ice. After sonication, 1 mL of ice-cold 10 μM ThT was added to the $A\beta_{42}$ solution and mixed by pipetting up and down 4 times, and then quickly transferred to the 15 mL conical tube containing the remaining 4.94 mL of ice-cold 10 μM ThT to create the 3.33 μM $A\beta_{42}$ solution. The $A\beta_{42}$ solution was then dumped into a sterile 25 mL reagent reservoir (ThermoFisher, catalog# 8093-11) and a 12-channel pipette was used to transfer 36 μL of the $A\beta_{42}$ solution to rows B-D.

**Reading the ThT assay plate.** The ThT assay plate was sealed with a clear adhesive plate seal (Axygen, catalog# PCR-SP) and was immediately inserted into a ThermoFisher Scientific Varioskan Lux plate reader. Fluorescence measurements of each well in rows C–H were acquired every 2 minutes over a 5 hour period with the following parameters:

| | |
|---|---|
| excitation: 440 nm | emission: 485 nm |
| measurement time: 1000 ms | optics: bottom read |
| excitation bandwidth: 12 nm | |

The data were plotted in GraphPad Prism.

#### Transmission Electron Microscopy

A 0.02 μmol aliquot of $A\beta_{42}$ was removed from the -80 °C freezer and allowed to equilibrate to room temperature. During this time, 6.78 μL of a 1.18 mg/ml stock solution of $pAb_{2AT-L}$ was diluted into 60 μL of PBS at pH 7.4 (10 mM $Na_2HPO_4$, 1.8 mM $KH_2PO_4$, 137 mM NaCl, 2.7 mM KCl) to yield a 888.89 nM solution of $pAb_{2AT-L}$. 27 μL of the 888.89 nM solution of $pAb_{2AT-L}$ was added to two low-binding microcentrifuge tubes. A third tube was treated with 27 μL of just PBS. The 0.02 μmol aliquot of $A\beta_{42}$ was then dissolved in 666.7 μL of PBS to yield a 30 μM solution. 3 μL of this solution was added to the microcentrifuge tube containing 27 μL of just PBS to yield a 3 μM solution. Another 3 μL of 30 μM $A\beta_{42}$ was added to one of the microcentrifuge tubes containing 27 μL of an 888.89 nM solution of $pAb_{2AT-L}$ to yield an 800 nM solution of $pAb_{2AT-L}$ in 3 μM $A\beta_{42}$. 3 μL of just PBS was added to the other tube containing 27 μL of an 888.89 nM solution of $pAb_{2AT-L}$ to yield an 800 nM solution of $pAb_{2AT-L}$. Samples were incubated for four hours at room temperature without shaking to elicit fibril formation.

200-mesh formvar/carbon-coated copper grid carbon-copper grids purchased from Electron Microscopy Sciences (catalog #FCF200-Cu-50) and carbon discharged under low vacuum by a Leica Sputter Coater ACE200 to increase the hydrophilicity of the grids. 5uL of sample was deposited on each grid and left to dry for 10 minutes. Afterwards, the grids were gently wicked and allowed to dry for another 5 minutes. The grids were then washed with 5 μL of nanopure water, wicked, and treated with 5 μL of two percent uranyl acetate for 2 minutes for negative staining. Afterwards, the grids were gently wicked and allowed to dry for another 5 minutes. Finally, the grids were washed with 5 μL of nanopure water and left to dry for 10 minutes before imaging. Samples were transferred to a JEOL 2100F TEM and imaged using a Schottky type field emission gun operating at 200kV. Images were recorded using a Gatan OneView CMOS camera at 4k x 4k resolution.

### References and Notes


1. Hoyer W, Grönwall C, Jonsson A, Ståhl S, Härd T. Stabilization of a β-Hairpin in Monomeric Alzheimer's Amyloid-β Peptide Inhibits Amyloid Formation. *Proceedings of the National Academy of Sciences* **2008**, *105* (13), 5099–5104. doi: 10.1073/pnas.0711731105.
2. Spencer RK, Li H, Nowick JS. X-ray Crystallographic Structures of Trimers and Higher-Order Oligomeric Assemblies of a Peptide Derived from $A\beta_{17–36}$ *Journal of the American Chemical Society* 2014, 136, 15, 5595–5598. doi: 10.1021/ja5017409
3. Kreutzer AG, Hamza IL, Spencer RK, Nowick JS. X-Ray Crystallographic Structures of a Trimer, Dodecamer, and Annular Pore Formed by an $A\beta_{17–36}$ β-Hairpin. *Journal of the American Chemical Society* **2016**, *138* (13), 4634–4642. doi: 10.1021/jacs.6b01332.
4. Haerianardakani S, Kreutzer AG, Salveson PJ, Samdin TD, Guaglianone GE, Nowick JS. Phenylalanine Mutation to Cyclohexylalanine Facilitates Triangular Trimer Formation by

β-Hairpins Derived from Aβ. *Journal of the American Chemical Society* **2020**, *142* (49), 20708–20716. doi: 10.1021/jacs.0c09281.

5. Ruttenberg SM, Kreutzer AG, Truex NL, Nowick JS. β-Hairpin Alignment Alters Oligomer Formation in Aβ-Derived Peptides. *Biochemistry* **2024**. doi: 10.1021/acs.biochem.3c00526.
6. Kreutzer AG, Parrocha CM, Haerianardakani S, Guaglianone G, Nguyen J, Diab MN, et al. Antibodies Raised against an Aβ Oligomer Mimic Recognize Pathological Features in Alzheimer's Disease and Associated Amyloid-Disease Brain Tissue. *ACS Central Science* **2023**. doi: 10.1021/acscentsci.3c00592.
7. Fernandopulle MS, Prestil R, Grunseich C, Wang C, Gan L, Ward ME. Transcription Factor-Mediated Differentiation of Human IPSCs into Neurons. *Current Protocols in Cell Biology* **2018**, *79* (1), e51. doi: 10.1002/cpcb.51.
